# Interdependence of EGFR with PTPs on juxtaposed membranes generates a growth factor sensing and responding network

**DOI:** 10.1101/309781

**Authors:** Angel Stanoev, Amit Mhamane, Klaus C. Schuermann, Hernán E. Grecco, Wayne Stallaert, Martin Baumdick, Yannick Brüggemann, Maitreyi S. Joshi, Pedro Roda-Navarro, Sven Fengler, Rabea Stockert, Lisaweta Roßmannek, Jutta Luig, Aneta Koseska, Philippe I. H. Bastiaens

## Abstract

The proto-oncogenic epidermal growth factor receptor (EGFR) is a tyrosine kinase whose sensitivity to growth factors (GFs) and signal duration determines cellular behavior. We resolve how EGFR’s response to epidermal growth factor (EGF) originates from dynamically established recursive interactions with spatially organized protein tyrosine phosphatases (PTPs). Opposed genetic PTP perturbations enabled identification of receptor-like PTPRG/J at the plasma membrane (PM) and endoplasmic reticulum (ER) associated PTPN2 as the major EGFR dephosphorylating activities. Imaging spatial-temporal PTP reactivity then revealed that vesicular trafficking establishes a spatially-distributed negative feedback with PTPN2 that determines signal duration, whereas single cell dose-response analysis uncovered a ROS-mediated toggle switch between autocatalytically activated monomeric EGFR and the tumor suppressor PTPRG that governs EGFR’s sensitivity to EGF. Vesicular recycling of monomeric EGFR unifies the interactions with these PTPs on juxtaposed membranes, dynamically generating a network architecture that can sense and respond to time varying growth factor signals.

## Introduction

Cells use cell surface receptors such a EGFR not only to sense the presence of extracellular growth factors but also to interpret the complex dynamic GF patterns that can lead to diverse, functionally opposed cellular responses including proliferation, survival, apoptosis, differentiation and migration (Yarden and Sliwkowski 2001). Collective molecular EGFR activity dynamics is thereby the first layer that translates the information encoded in time varying extracellular GF patterns into a cellular outcome. Such a system must have two essential characteristics: sensitivity to time varying GF inputs and capability to transform these inputs into an intracellular spatial temporal activity pattern. EGFR phosphorylation dynamics that reflects these complex features however cannot be accounted for using the classical view how information about GFs is written and erased by kinases and phosphatases, since this only refers to sensing presence or absence of GFs. In particular, canonical EGFR activation by GFs relies on dimerization and allosteric activation of its intrinsic kinase activity, which results in the phosphorylation of tyrosine residues on the C-terminal receptor tail (Arkhipov et al., 2013; Kovacs et al., 2015; Schlessinger, 2002) that serve as docking sites for SH2-or PTB-containing signal transducing proteins (Wagner et al., 2013). In this view PTPs serve as mere erasers of the signal that is written by the intrinsic kinase-dependent phosphorylation of the receptor (Lim and Pawson, 2010). Complex response dynamics can however emerge from recursive interactions with PTPs in combination with autocatalytic receptor activation that gives rise to specific network modules (Baumdick et al., 2015; Grecco et al., 2011; Koseska and Bastiaens, 2017; Reynolds et al., 2003; Schmick and Bastiaens, 2014; Tischer and Bastiaens, 2003). Large scale studies based on enzymatic assays of purified PTPs (Barr et al., 2009), membrane two-hybrid assays (Yao et al., 2017) and biochemical assays on cell extracts after siRNA knockdown (Tarcic et al., 2009) have identified novel or confirmed known PTPs that dephosphorylate EGFR (Liu and Chernoff, 1997; Tiganis et al., 1998; Yuan et al., 2010). However, which PTPs determine the collective phosphorylation dynamics of EGFR, leading to sensing of growth factor patterns and a subsequent cellular response remains unknown.

To first identify the major PTPs that dephosphorylate EGFR we used cell array fluorescence lifetime imaging microscopy (CA-FLIM) screening (Grecco et al., 2010) in combination with quantifiable genetic perturbations. We then addressed how these PTPs with distinct localizations (Tonks, 2006) couple to EGFR and spatially regulate the EGFR phosphorylation response to time varying growth factor inputs. By decomposing EGFR phosphorylation in space and time, we uncovered that a spatially distributed EGFR-PTPN2 negative feedback regulates the temporal EGFR phosphorylation response, whereas single-cell dose-response analysis enabled the derivation of the underlying PTP-EGFR network architecture that dictates sensitivity to EGF dose. This revealed that a reactive oxygen species-mediated toggle switch between EGFR and the tumor suppressor PTPRG (Kwok et al., 2015) is central to determine EGFR’s sensitivity to EGF dose. The interdependence of EGFR and PTPRG is further modulated by negative regulation by PTPRJ at the PM. Vesicular dynamics in the form of constitutive vesicular recycling repopulates the PM with monomeric receptors to enable autocatalysis but also unifies recursive interactions between EGFR and PTPRs at the PM with PTPN2 on the ER in a network with specific topology. This identified EGFR-PTP system enables sensing of time-varying GF stimuli.

### Vesicular recycling of ligandless EGFR determines phosphorylation dynamics

To investigate how PTPs determine EGFR’s response to growth factors, we first assessed how EGFR’s phosphorylation dynamics relates to EGF dose and its vesicular trafficking. To observe EGFR phosphorylation dynamics at physiological expression, fluorescently tagged EGFR-mTFP was ectopically expressed in breast cancer-derived MCF7 cells with low endogenous EGFR expression (~ 10^3^/cell (Charafe-Jauffret et al., 2005), Figure S1A), to a level that fell within the endogenous EGFR expression range of the related MCF10A cells (determined by EGF-Alexa647 binding, Figure 1A). EGFR-mTFP expressing MCF7 cells exhibited equivalent EGFR phosphorylation-(Y_1068_-Grb2 binding site (Okutani et al., 1994)) and Akt activation dynamics to MCF10A cells in response to 200ng/ml of sustained (S-EGF) as well as 5 min pulsed (5P-EGF) EGF-Alexa647 stimulus (Figure 1B). This shows that EGFR-mTFP expressing MCF7 cells exhibit physiological EGFR response properties.

**Figure 1.**
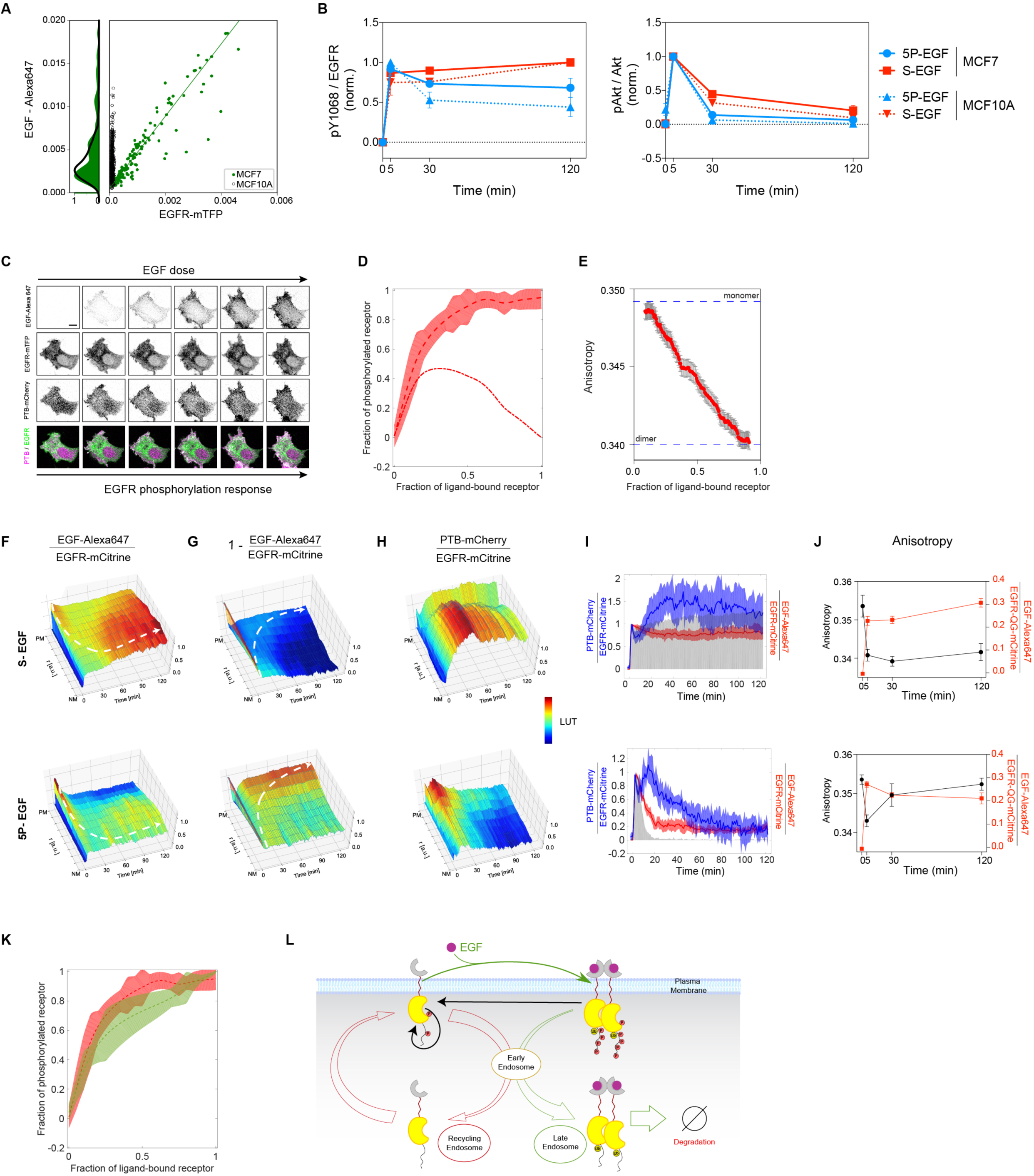
EGFR phosphorylation and vesicular dynamics. **(A)** Quantifying ectopic EGFR-mTFP expression in MCF7 cells. Average EGF-Alexa647 vs. EGFR-mTFP fluorescence in single MCF7 (green) or MCF10A cells without EGFR-mTFP (black). Histograms reflect that levels of EGF-Alexa647 binding to MCF7 with ectopic EGFR-mTFP expression (green) and MCF10A with endogenous EGFR (black) are similar. **(B)** EGFR Y_1068_ phosphorylation (left) and Akt phosphorylation (right) in MCF7 cells ectopically expressing EGFR-mTFP (solid lines) and endogenous EGFR in MCF10A cells (dashed lines), following 5min pulsed (5P-EGF, 200ng/ml, blue) or sustained EGF stimulation (S-EGF, 200ng/ml, red), as determined by in-cell Western assay (N=3). Data are normalized to the maximum response in each respective condition (means ± SEM, N=3). **(C)** Representative fluorescence image series of EGF-Alexa647, EGFR-mTFP, PTB–mCherry and PTB-mCherry(magenta)/EGFR-mTFP(green) overlay from singlecell dose-response experiment. Cells were stimulated every ~1.5min with increasing EGF-Alexa647 doses (2.5-600ng/ml). Scale bar: 20μm. (**D**) Fraction of phosphorylated vs. ligand-bound EGFR-mTFP (n=21, N=2; Figure S1B-D). Dashed lines: moving averages from single-cells; shaded bounds: standard deviations; dash-dotted lines: estimated contribution of ligandless to the fraction of phosphorylated EGFR. (**E**) Livecell fluorescence anisotropy microscopy measurements of EGFR-QG-mCitrine dimerization state as a function of the fraction of ligand-bound receptor (mean ± SEM, n=30, N=3, Figure S1F,G). **(F-H)** Average spatial-temporal maps (STMs) of the estimated fraction of ligand-bound EGFR (**(F),** EGF-Alexa647/EGFR-mCitrine), ligandless EGFR (**(G),** 1-[EGF-Alexa647/EGFR-mCitrine]) and the fraction of phosphorylated EGFR-mTFP estimated by PTB-mCherry translocation (**(H)**, PTB-mCherry/EGFR-mCitrine). Data was acquired at 1min-intervals in live MCF7 cells following 200ng/ml S-EGF (top, n=16, N=3; Figure S1I,J) or 5P-EGF (n=14, N=2; Figure S1I,J) stimulation. White dotted lines: trajectories representing the change in distribution of ligand-bound (F) and ligandless (G) EGFR. **(I)** The respective plasma membrane fractions of ligand-bound (EGF-Alexa647/EGFR-mCitrine, red) and phosphorylated EGFR (PTB-mCherry/EGFR-mCitrine, blue) derived from **(F)** and **(H)** (median ± AMD). Extracellular EGF-Alexa647 are shown in grey. **(J)** Dimerization state (black) and the fraction of ligand-bound EGFR-QG-mCitrine (red) at the PM for live cells following 200ng/ml S-EGF following 200ng/ml S-EGF (top, n=5, N=3) or 5P-EGF (bottom, n=5, N=3) stimulation (means ± SEM). **(K)** The dose-response of EGFR-mTFP phosphorylation (red, control) is significantly altered upon ectopic Rab11a^S25N^ expression (green, *p*=0.02, n=12, N=3). Lines same as in **(D)**. **(L)** Scheme of EGFR trafficking dynamics: ligandless EGFR recycles via early (EE) and recycling endosomes (RE) to the PM (red arrows) whereas upon EGF binding (thin green arrow), ubiquitinated EGF-EGFRub unidirectionally traffickes via the early-to the late endosomes (LE, green arrow) to be degraded (Ø). Causal links are denoted with solid black lines.

To first assess the sensitivity of EGFR phosphorylation response to EGF binding, we performed single cell dose-response experiments with fluorescent EGF-Alexa647 (Figure 1C). To deconvolute EGF binding kinetics from EGFR’s response, we directly related the fraction of liganded receptors to EGFR phosphorylation, which is not possible by analytical biochemical approaches on cell extracts. The fraction of liganded EGFR-mTFP at the PM was determined by EGF-Alexa647/EGFR-mTFP, and EGFR-mTFP phosphorylation was measured by the rapid translocation of mCherry-tagged phosphotyrosine-binding domain (PTB-mCherry, Figure S1E) to the phosphorylated tyrosines 1086/1148 of EGFR at the PM (Offterdinger et al., 2004) (Figure S1B-S1E, STAR Methods).

The observed steep EGFR phosphorylation response (Figure 1D, S1D) showed that the largest fraction of phosphorylated receptors at low EGF doses are ligandless (dash-dotted line in Figure 1D, STAR Methods), pointing to an amplification of ligandless EGFR phosphorylation that contributes to this steepness. The high fraction of phosphorylated EGFR at low fraction of liganded receptors additionally indicates that liganded EGFR triggers the phosphorylation amplification on ligandless EGFR. Measuring the dimerization state of EGFR as function of EGF doses by homo-FRET detection with fluorescence anisotropy microscopy on a fully active EGFR-QG-mCitrine construct (Baumdick et al., 2015), showed that the fraction of ligand bound receptors corresponds to the fraction of dimerized EGFR (Figure 1E, S1F-G). From this, it can be deduced that phosphorylated ligandless EGFR is monomeric.

Given the contribution of ligandless monomers to the sensitivity of EGFR activation, we investigated how vesicular dynamics relates to EGFR phosphorylation by exposing cells to both sustained (S-EGF) as well as pulsed (5P-EGF) stimulation. For this, EGFR-mTFP, EGF-Alexa647 as well as PTB-mCherry fluorescence distributions were monitored by live cell confocal microscopic imaging, and receptor self-association was monitored in a separate experiment by fluorescence anisotropy microscopy on EGFR-QG-mCitrine. The molecular quantities of ligandless EGFR fraction at each pixel was calculated from: 1-[EGF-Alexa647/EGFR-mTFP], and EGFR phosphorylation from: PTB-mCherry/EGFR-mTFP. The radial symmetry in receptor trafficking from the PM to the nuclear membrane (NM) enabled dimensionality reduction of the Cartesian variables (x, y) to normalized radial variable (r, Figure S1H), which allowed us to reconstruct average 3D spatial-temporal maps from multiple cells (STMs in Figure 1F-H, S1I-J, STAR Methods).

Upon sustained EGF stimulation, liganded dimers (Figure 1F top, EGF-Alexa647/EGFR-mTFP) at the PM were activated (Figure 1H,1I top, PTB-mCherry/EGFR-mTFP), endocytosed, and unidirectionally trafficked towards the perinuclear area in the course of 2h, where they were inactivated by dephosphorylation (Figure 1H, S1I, Movie S1). Retrograde trafficking of ligandless receptors from the perinuclear recycling endosome (RE) to the PM (Baumdick et al., 2015) was also observed following S-EGF stimulation (Figure 1G, top), where they immediately bound EGF. This was reflected in the continuous high fraction of dimers at the PM, as measured by the anisotropy of EGFR-QG-mCitrine (Figure 1J, top).

To next investigate if receptors can auto-phosphorylate after a stimulus is removed, we exposed cells to a 5min pulse of EGF (5P-EGF) and spatially resolved EGFR’s phosphorylation dynamics over 2h. During the 5 min EGF pulse, receptors bound EGF and were depleted from the PM to accumulate in the perinuclear area where they were dephosphorylated (Figure 1F, H, bottom). However, in the time after the 5P-EGF pulse, ligandless receptors rapidly recycled to the PM (t_1/2_~2-4 min, STAR Methods; Figures 1J bottom, S1K; Movie S2) where they were re-phosphorylated in the absence of ligand, exhibiting their maximal phosphorylation at ~15 min after 5P-EGF to then slowly decay to a dephosphorylated state (Figure 1I, bottom). Fluorescence anisotropy measurements of EGFR-QG-mCitrine at the PM showed that the recycled EGFR was monomeric (Figure 1J, bottom). In accordance with this, the blocking of vesicular recycling by ectopic expression of dominant negative Rab11^DN^ mutant (Konitsiotis et al., 2017) led to a significant decrease in the steepness of the EGFR phosphorylation response (Figure 1K).

These experiments thus show that ligandless and liganded EGFR exhibit distinct vesicular and phosphorylation dynamics that can be distinguished by 5P-EGF stimulus (Figure 1L). Upon ligand binding, ligandless EGFR is transformed to dimeric EGFR (green arrow, Figure 1L). The dimers can in turn activate auto-phosphorylation on remaining or recycling monomeric EGFR (black arrow, Figure 1L), thereby amplifying the response. In contrast to the recycling ligandless monomeric EGFR which can additionally get reactivated by autocatalysis at the PM (Baumdick et al., 2015), liganded dimeric EGFR uni-directionally traffics to late endosomes. This indicates that a continuously maintained PM-fraction of EGFR monomers allows for sensing of time-varying GF stimuli.

### The major PTPs that dephosphorylate EGFR are on juxtaposed membranes

To investigate how PTPs regulate EGFR phosphorylation in this vesicular dynamic system, we identified which PTPs have the strongest dephosphorylating activity on EGFR. We used opposed genetic perturbations, where siRNA mediated knockdown allowed the identification of PTPs that have non-redundant EGFR dephosphorylating activity, whereas ectopic PTPx-mCitrine expression enabled the derivation of their specific enzymatic activity on phosphorylated EGFR. The change in EGFR phosphorylation in response to these opposed genetic perturbations was measured by determining the change in Förster Resonance Energy Transfer (FRET) that occurs upon binding of an anti-phosphotyrosine antibody tagged with Cy3.5 (PY72-Cy3.5) to phosphorylated EGFR-FP (Wouters and Bastiaens, 1999). FRET was measured via the decrease in fluorescence lifetime of EGFR-FP in single cells using CA-FLIM and the fraction of phosphorylated EGFR-FP (α) was quantified using global analysis (Grecco et al., 2010) (Figure S2A-B). The effect of the genetic PTP perturbations on EGFR phosphorylation was then determined by the phosphorylation fold change (PFC): *PFC_α_* = *α_PTP_*/*α_ctr_*.

CA-FLIM screening of 55 PTPs that are expressed in MCF7 cells (Figure S1A; Tables S2, S3) and well represent the four PTP families (Alonso et al., 2004), showed that 39 significantly affected EGFR phosphorylation (*PFC_α_*) after 5min of EGF stimulation. However, only 5 PTPs increased EGFR phosphorylation upon knockdown (*PFC_α_*-siRNA) and decreased it upon ectopic PTP_X_-mCitrine expression (*PFC_α_* – *cDNA*), identifying them as non-redundant negative regulators of EGFR phosphorylation (Figure 2A, red dots in quadrant 1, diameter proportional to mRNA expression in MCF7 cells). These were the ER-bound PTPN2 (Lorenzen et al., 1995) and the receptor-like PTPRG/J/A (Andersen et al., 2001; Barr et al., 2009) belonging to the family of classical PTPs, as well as the dual-specificity phosphatase DUSP3. Additionally, the lowly expressed DUSP7 and DUSP10 were identified as positive regulators with both genetic perturbations (Figure 2A, red dots in quadrant 3). These are necessarily indirect effectors, implicating that the expression level of auxiliary proteins does not limit their positive regulation of EGFR phosphorylation.

**Figure 2.**
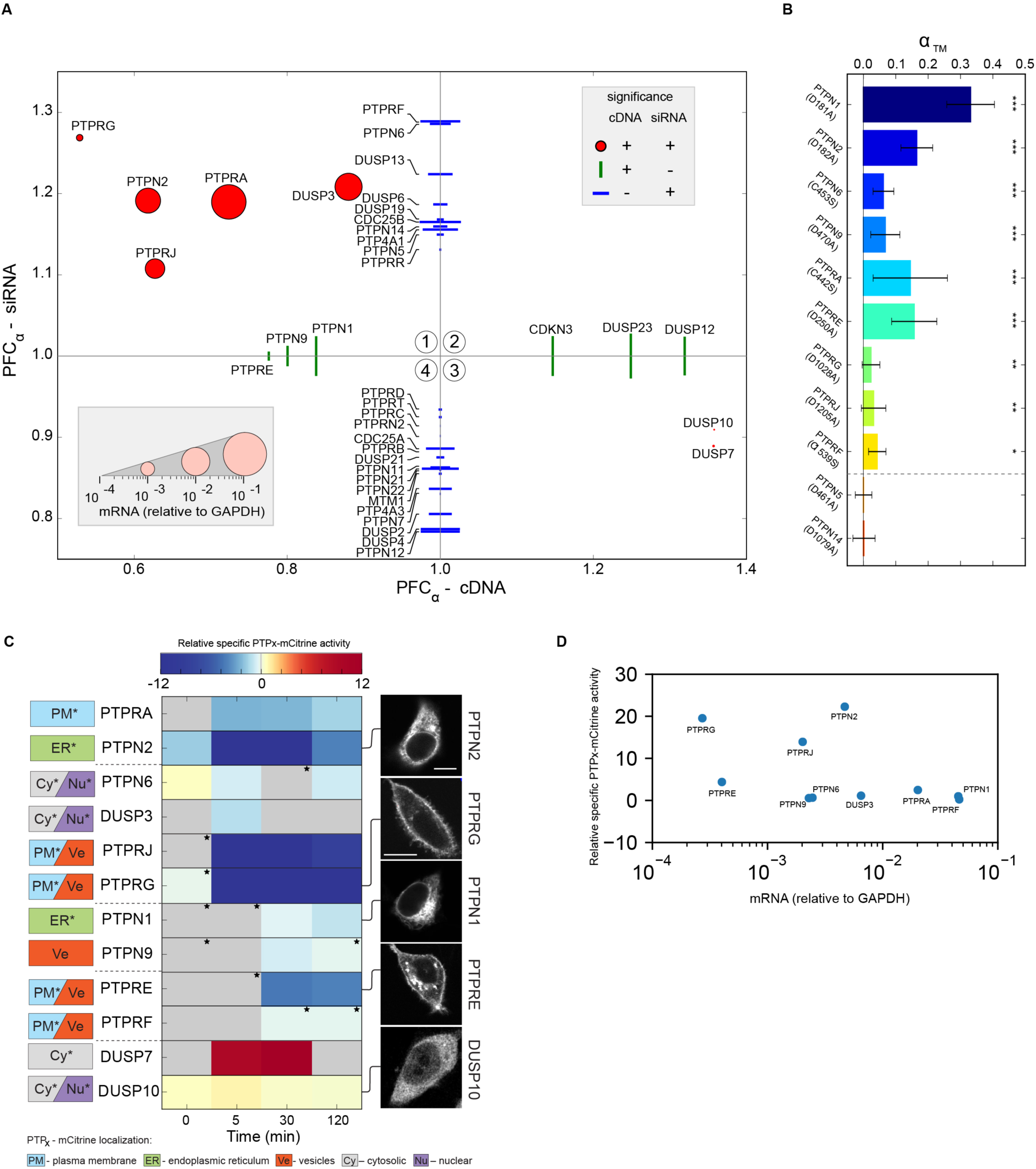
In situ EGFR phosphatome identification. **(A)** Scatter plot of median EGFR phosphorylation fold-changes (*PFC*_*α*_ = ^*α*_*PTP*_^/*α*_*ctr*_, n~150 cells per condition) upon siRNA-knockdown (*PFC*_*α*_-siRNA) and ectopic PTP_x_-mCitrine expression (*PFC*_α_-cDNA), 5min after 200ng/ml EGF stimulation. Significant *PFC*_*α*_ upon both (red dots) or only one perturbation (green/blue lines, p<0.05) are shown. Marker length is scaled to relative PTP_X_-mRNA expression in MCF7 cells (legend:inset and Figure S1A). **(B)** Average fraction of EGFR-mTFP interacting with catalytically impaired PTPX-mCitrine trapping mutants (aTM ± SD, n=15-20 cells, * p<0.05, ** p<0.01 and *** p<0.001) following 200 ng/ml 5P-EGF. **(C)** Relative specific PTPX-mCitrine activities (middle) prior to and 5, 30 and 120min after 200ng/ml 5P-EGF stimulation. *: Weak linear dependencies (Figure S2D). Subcellular localization of PTPX-mCitrine (left boxes, *: additionally curated localization from LOCATE/UniProt database) and exemplary fluorescence images (right, scale bar: 10 μm) **(D)** Relative specific PTPX-mCitrine activities versus the corresponding mRNA expression in MCF7 cells.

Most of the remaining 32 PTPs affected EGFR phosphorylation only upon knockdown (*PFC*_*α*_*-siRNA*, blue lines, Figure 2A), whereas 6 have an effect only upon ectopic expression (*PFC*_*α*_ – *cDNA*, green lines, Figure 2A). The majority of these PTPs fell on the right of the cDNA and below the siRNA axis, and are therefore indirect positive regulators of EGFR phosphorylation. On the other hand, the effect of the negative regulators that manifests only upon a single genetic perturbation reflects either redundancy in case of ectopic expression, or PTPs whose activity depends on and is limited by the amount of phosphorylated EGFR in case of knockdown.

FLIM-FRET measurements of the interaction of EGFR-mTFP with fluorescent, catalytically impaired PTP trapping variants (Flint et al., 1997), showed that the four classical non-redundant negative regulators (PTPN2, PTPRG/J/A) (Figure 2B, Table S3) and the redundant negative regulators (PTPN1/9, PTPRE; identified upon ectopic expression), as well as the strongest negative regulators identified upon knockdown (PTPN6, PTPRF) directly dephosphorylate EGFR. On the other hand, interaction with EGFR-mTFP was not observed with the trapping variants of indirect negative regulators (PTPN5, PTPN14) (Belle et al., 2015; José et al., 2003).

In order to determine which of the identified PTPs exert the strongest dephosphorylating activity on EGFR, we used cell-to-cell variance in PTPx-mCitrine expression to determine the specific activity of each of these PTPs towards EGFR-mTFP. For this, we measured EGFR-mTFP phosphorylation (*α*_t_, Figure S2C) and ectopic PTP_X_-mCitrine expression in individual cells upon 5P-EGF to generate scatter plots of the fraction of phosphorylated EGFR (α) vs. PTP_X_-mCitrine fluorescence for a given time point (Figure S2D). To obtain the specific activities, the scatter plots were parameterized by an exponential fit. This showed that three of the non-redundant negative regulators identified from the opposed perturbations (PTPN2 and PTPRG/J) exhibited the strongest dephosphorylating activity towards EGFR-mTFP (Figure 2D) that extended over the fulltime range after EGF stimulation (Figure 2C). These three strongest regulators have juxtaposed subcellular localizations of cytoplasmic ER (PTPN2) and the PM (PTPRG/J) (Figure 2C), where the PTPRs exhibited one to two orders of magnitude lower mRNA expression than PTPN2 (Figure 2D). In contrast, the highly expressed soluble DUSP3 and PM-localized PTPRA were profiled as only weak or modest regulators of EGFR phosphorylation, respectively.

### ER-PTPs and PTPRs dynamically shape EGFR’s phosphorylation response in space

To determine how the juxtaposed PTPs shape EGFR phosphorylation in space, the effect of opposed genetic PTP_×_ perturbations on EGFR-Y_1068_ phosphorylation after 5P-EGF was imaged in many individual cells by immunofluorescence using a specific pY_1068_ antibody (Figure S3A-B). From these images, we reconstructed 3D spatial-temporal maps of the average fraction of phosphorylated EGFR-mTFP (pY_1068_–Alexa568/EGFR-mTFP, Figure 3A, STAR Methods) at 0, 5, 30 and 120 min following 5P-EGF stimulation. To map where the PTPx dephosphorylates EGFR, the genetic perturbation effects were quantified by the phosphorylation fold change (PFC) relative to control (ctr) defined by:

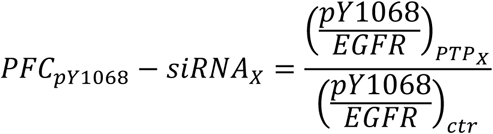

for knockdown of a PTPX (Figure S3C), and:

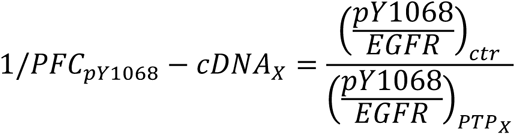

for ectopic PTP_x_-mCitrine expression. Average PFCs of many cells were accumulated using the same dimensionality reduction and distance normalization as in Figure 1F-H. Mathematical modeling of the phosphorylation/dephosphorylation cycle showed that the experimentally derived 1/*PFC*_*pY*1068_ – *cDNA*_*X*_ approximates the local specific dephosphorylating activity of an ectopically expressed PTPx-mCitrine relative to the local kinase activity of EGFR1/*PFC*_*pY*1068_-*cDNA_X_*≈*k_ptpx_[PTP^X^]*/*k_EGFR_*; STAR Methods). To avoid loss of spatial information on these activities due to PTP_X_-mCitrine overexpression-induced saturation, we only analyzed cells where pY_1068_/EGFR depended linearly on PTPx-mCitrine (Figure S3D).

**Figure 3.**
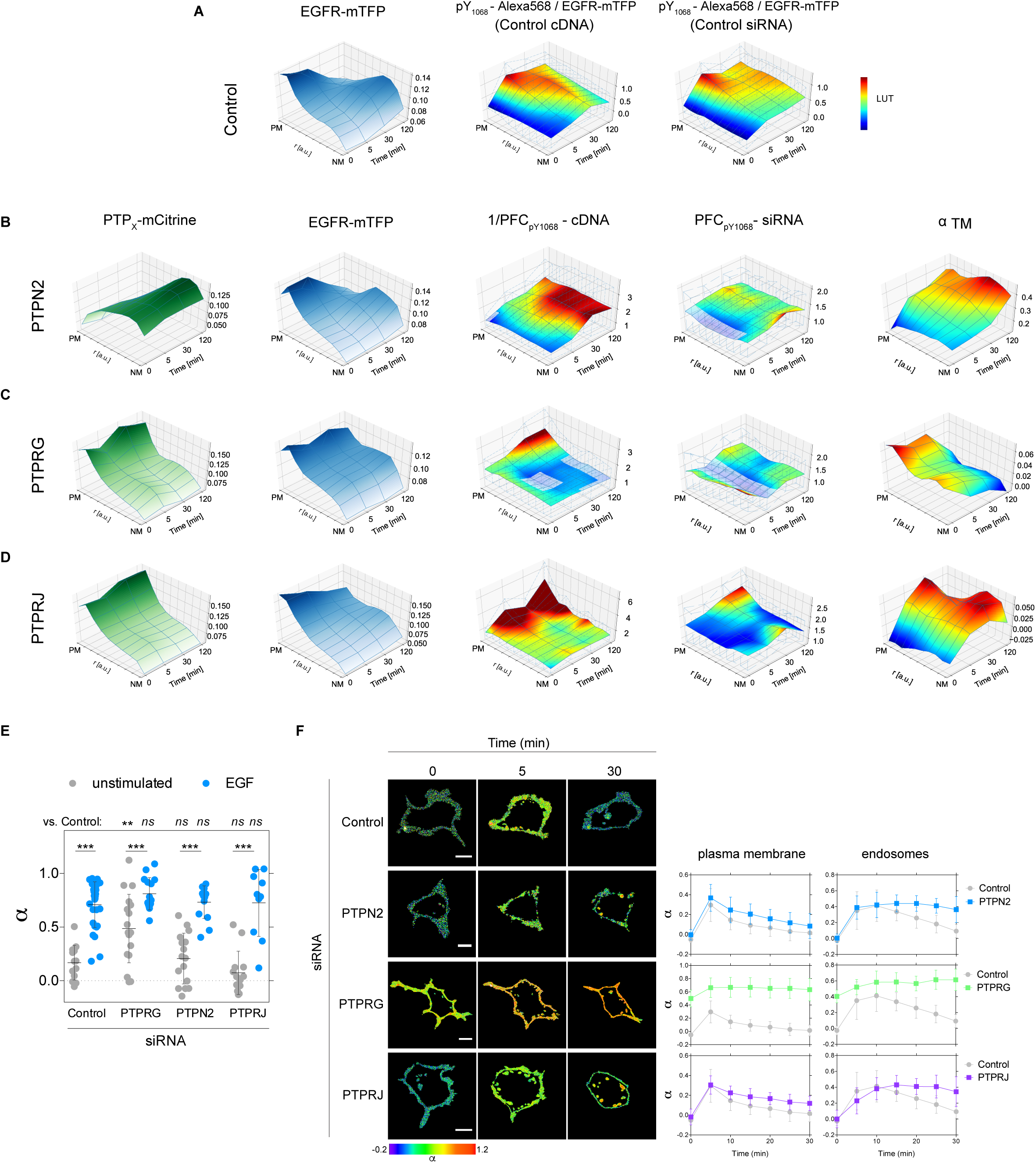
Spatial-temporal regulation of EGFR phosphorylation by ER-bound and PTPRs. **(A)** Spatial-temporal maps (STMs) depicting EGFR-mTFP fluorescence (left) and pY_1068_ phosphorylation (middle) in control cells (n~90 cells per time point, N=6 experiments) and following transfection with non-targeting siRNA pool (right, n~60, N=4). **(B)** Columns 1-3: effect of PTPN2-mCitrine expression (Column 1) on STMs of EGFR-mTFP localization (Column 2) and phosphorylation fold-change (1/PFC_pY1068_-cDNA, Column 3), which reflects the relative PTPN2-mCitrine reactivity towards pY_1068_ (n~60, N=3). Column 4: effect of siRNA-mediated PTPN2 knockdown on EGFR-mTFP phosphorylation fold change (PFC_pY1068_-siRNA, n~45, N=3). Column 5: STM of fraction of EGFR-mTFP interacting with PTPN2^C216S^-mCitrine trapping mutant (α_™_, n=15-30). **(C)** STMs of the same quantities as in **(B)** upon PTPRG-mCitrine expression (n~60, N=3; α_™_ PTPRG^C1060S^-mCitrine n=15-30). **(D)** STMs of the same quantities as in **(B)** upon PTPRJ-mCitrine expression (n~40, N=2; α_™_ PTPRJ^D1205A^-mCitrine, n~30). In **(A-D)**, cells were stimulated with 200ng/ml 5P-EGF; transparent areas: non-significant PFCs, p >0.05. **(E)** Effect of siRNA-mediated knockdown of PTPRG, PTPN2 and PTPRJ on the fraction of phosphorylated EGFR (α) in single MCF7 cells expressing EGFR-mCitrine (donor) and PTB-mCherry (acceptor). FLIM measurements were made prior to (grey) and 2min following saturating 320ng/ml EGF-Alexa647 stimulation (blue). amean ± SD for Control: n=14 (grey), n=17 (blue); PTPRG: n=15 (grey), n=11 (blue); PTPN2: n=9 (grey), n=8 (blue); PTPRJ: n=6 (grey), n=6 (blue); is shown. N=1-2. ** p=0.0018 and *** p<0.001. **(F)** Time-lapse measurements of the fraction of phosphorylated EGFR (as above) in single MCF7 cells prior to and every 5 minutes after 200 ng/ml 5P-EGF stimulation for a total of 30 min. Representative α images (left) and corresponding quantifications (right) for control (n=4), PTPN2 (n=5), PTPRG (n=5) and PTPRJ (n=4) knockdowns (N=3). Scale bar: 10 μm.

The STM of PTPN2-mCitrine fluorescence shows that PTPN2 concentration steadily declines from the perinuclear area towards the cell periphery, invariant over time (Figure 3B), whereas the profile of fluorescent EGFR-mTFP reflected the typical vesicular dynamic behavior of internalization and recycling after 5P-EGF. Thus, as phosphorylated EGFR traffics from the PM via early endosomes (EE) to late endosomes (LE) or REs along this increasing PTPN2-concentration, it is progressively dephosphorylated on p_Y1068_ (Figure 3B, PFCs). Both *PFC_pYW68_-siRNA_pTPN2_* and 1/*PFC_pyw68_-cDNA_PTPN2_* additionally showed an increasing dephosphorylating activity of PTPN2 with time at the cell periphery, revealing that a fraction of ER-bound PTPN2 can reach the PM (Lorenzen et al., 1995) to dephosphorylate EGFR-pY_1068_. This was corroborated by the interaction profile of EGFR-mTFP with the trapping PTPN2^C216S^-mCitrine variant (Tiganis et al., 1998), which increased both towards the perinuclear- and the peripheral cytoplasm over time (Figure 3B, STM-α_™_).

PTPRG-mCitrine displayed strong fluorescence at the cell periphery that abruptly declined in the cytoplasm, but in contrast to PTPN2 exhibited dynamic redistribution after stimulation (Figure 3C, STM PTP_x_-mCitrine). This redistribution of PTPRG coincided with that of EGFR, initially internalizing in endosomes, to then traffic back and gradually increase at the PM. This points at a direct interaction of PTPRG and EGFR. The *PFC_pY1068_-siRNA_pTPRG_* showed an enhanced phosphorylation of EGFR in the absence of stimulus, indicating that PTPRG maintains EGFR monomers dephosphorylated. After 5P-EFG, both *PFC_pY1068_-siRNA_pTPRG_* and *PFC_pY1068_-cDNA_pTPRG_* revealed a steady increase in PTPRG activity at the PM over time (Figure 3C).

PTPRJ-mCitrine distribution did not coincide with that of EGFR, translocating from endosomes back to the PM late after 5P-EGF (Figure 3D). In stark contrast to PTPRG, the dephosphorylating activity of PTPRJ was low in the absence of stimulus and increased after 5P-EGF, following its observed redistribution to the PM. The differences in the interaction of EGFR-mTFP with the trapping variants of the two PTPRs (STM-α_™_, Figure 3C-D) reflect their differences in regulating EGFR dephosphorylation. Whereas the interaction of the PTPRG^C1060S^-mCitrine (Table S3) with EGFR-mTFP already occurred in the absence of stimulus (STM-α_™_ 0min, Figure 3C), the interaction with PTPRJ^D1205A^-mCitrine was apparent and increasing only after 5P-EGF (STM-α_™_, Figure 3D). This indicates that PTPRG preferentially dephosphorylates ligandless EGFR at the PM, corroborated by the strongly reduced PTPRG activity upon S-EGF stimulus when the majority of the receptor is liganded (Figure S3E). Immuno-precipitation experiments further confirmed that there is a preferential interaction of PTPRG-mCitrine with ligandless EGFR (activated due to PTP inhibition by H_2_O_2_ (Meng et al., 2002)) over liganded EGFR activated by EGF, whereas PTPRJ constitutively interacts with both species (Figure S3H). Interestingly, the STM-aTM of PTPRJ^D1205A^-mCitrine also showed an increase of interaction in the perinuclear cytoplasm after 5P-EGF, which is consistent with the *PFC*_*pY1068*_*-siRNA*_*PTPRJ*_ and indicates that an intracellular endosomal fraction of PTPRJ dephosphorylates endocytosed EGFR.

The more static spatial-temporal distribution of the other identified non-redundant receptor-like PTPRA-mCitrine did not coincide with that of EGFR (Figure S3F). Even more, its specific activity towards EGFR-pY_1068_ increased at intermediate and late times after EGF stimulation, following the interaction profile of EGFR-mTFP with the trapping PTPRA ^C442S^-mCitrine variant. This indicates that PTPRA suppresses autonomous activation of recycling ligandless receptors mostly at the PM late after stimulus (*PFC_pY1068_-siRNA_PTPRA_*, 1/*PFC_py1068_-cDNA_PTPRA_*, Figure S3F). siRNA-mediated knockdown of DUSP3 confirmed the low specific activity (Figure 2C) of this atypical phosphatase towards EGFR-p_Y1068_ (Figure S3G).

To further investigate how the three strongest PTPs affect EGFR phosphorylation dynamics, we measured time-lapse EGFR-mTFP phosphorylation response to 5P-EGF in living cells upon PTPx knockdown. EGFR phosphorylation was imaged via the interaction of PTB-mCherry with phosphorylated EGFR-mCitrine by FLIM and quantified by global analysis (Grecco et al., 2010) to obtain the average fraction of phosphorylated EGFR (α) at the PM and on endosomes (Figure 3E-F, STAR Methods). Since only PTPRG knock down resulted in elevated basal EGFR phosphorylation and its trapping variant already interacted with EGFR in the absence of stimulus (Figure 3C), we compared α upon knock-down of these three PTPs in the absence of EGF stimulus in single cells (Figure 3E). This clearly showed that PTPRG knock-down resulted in substantial pre-activated EGFR. The wide distribution of EGFR phosphorylation in this case likely reflects the variance in PTPRG knock-down level in each cell. Consistently, time-lapse FLIM of EGFR phosphorylation showed the already high EGFR phosphorylation on the PM and in endosomes in the absence of stimulus to only slightly increase to a plateau after 5P-EGF (Figure 3F). PTPRJ knock-down resulted in more sustained phosphorylation of EGFR monomers at the PM after 5P-EGF. Interestingly, a steady increase in the phosphorylation on endosomes could be observed that plateaued 15 min after 5P-EGF. This indicates that knock-down of PTPRJ increases the phosphorylation of recycling EGFR monomers. In contrast, PTPN2 knock-down only changed the amplitude of the response at the PM without affecting its profile, whereas activation of EGFR signaling from endosomes initially followed that at the PM but was then clearly sustained at later times.

These results are consistent with the PFCs (Figures 3B-D) and show that PTPN2 determines signal duration by dephosphorylating liganded EGFR during its vesicular trafficking, whereas PTPRG and PTPRJ dephosphorylate recycling ligandless EGFR. This suggests that PTPRG/J most likely have a functional role in determining the sensitivity of EGFR phosphorylation response to EGF.

### PTPRG and PTPRJ are central regulators of EGFR responsiveness to EGF dose

To understand how EGFR sensitivity to GFs is regulated by the distinct activity of PTPRG/J at the PM and PTPN2 on the ER, we determined EGFR-mTFP phosphorylation response to EGF dose upon PTPx knockdown. This was performed in single cells analogous to the experiments presented in Figure 1C-D. The pre-activation of EGFR phosphorylation upon PTPRG knockdown (Figure 3E) impedes EGFR responsiveness to EGF, and we therefore did not perform this experiment. Strikingly, PTPRJ knockdown induced a more switch-like EGFR phosphorylation response (Figure 4A bottom and S4E), whereas knockdown of PTPN2 significantly steepened the EGF-dose response (Figure 4A, middle). Consistent with its late function in suppressing autonomous activation of recycling receptors at the PM (Figure S3F), knock-down of PTPRA did not affect the EGF dose-EGFR phosphorylation response (Figure S4D).

**Figure 4.**
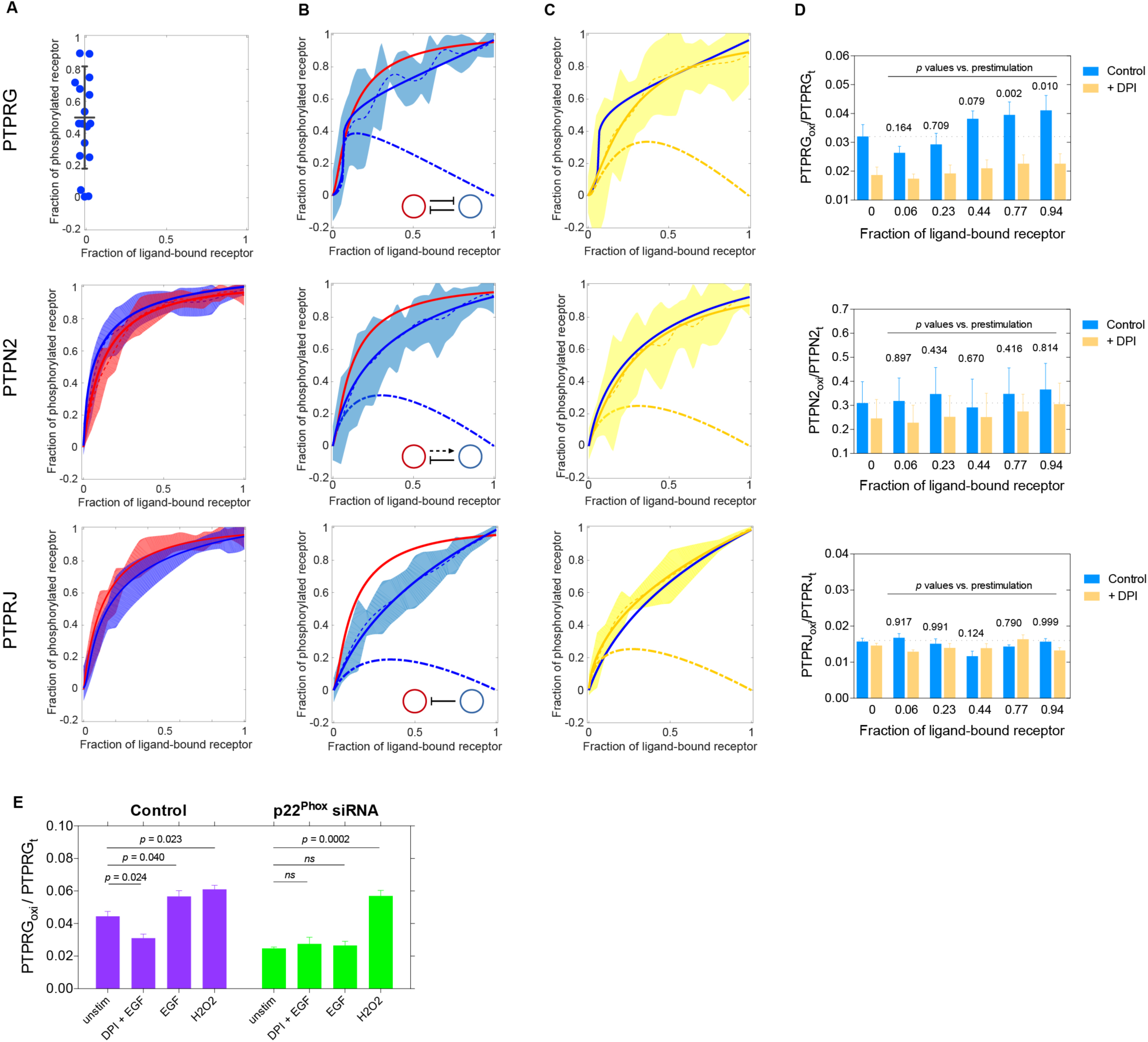
Differential regulation of EGFR responsiveness by ER-bound-and R-PTPs. **(A)** Averaged single-cell dose-response measurements following PTP_x_ knockondown. PTPRG knockdown results in preactivated EGFR phosphorylation (top, blue dots on the Y-axis). Dose-response of EGFR-mCitrine phosphorylation (red, n=21, N=4) is significantly altered upon siRNA-mediated PTPRJ knockdown (bottom, blue line, p=0.004; n=11, N=3) and less upon PTPN2 (middle, blue line, p=0.17; n=14, N=4) knockdown. Shaded bounds: identical as in Figure 1D. Solid lines: model-based fits to the phosphorylated EGFR fraction (STAR Methods, Figure S4A). **(B)** Dose-response of EGFR-mTFP phosphorylation (red) is significantly altered upon PTPRG-mCitrine coexpression (blue lines, p=0.027; n=28, N=4; top), PTPN2-mCitrine (blue lines n=34, N=5, p=0.001; middle), or PTPRJ-mCitrine co-expression (p=4*10-4; n=6, N=2; bottom). Solid lines: model-based fits to the phosphorylated EGFR fraction (STAR Methods, Figure S4A). Best fits are with the model shown in inset. **(C)** NOX-inhibition by DPI (10μM, 30min pre-incubation) significantly flattens dose-response of EGFR phosphorylation upon ectopic PTPRG-mCitrine (yellow lines, p=0.06, n=26, N=3, top), but has no effect upon PTPN2-mCitrine (p=0.19, n=45, N=3; middle) or PTPRJ-mCitrine expression (p=0.162, n=10, N=2; bottom). **(D)** Quantification of PTPRG-mCitrine (top), PTPN2-mCitrine (middle) and PTPRJ-mCitrine (bottom) catalytic cysteine oxidation for different EGF-Alexa647 doses (blue bars, N=4-7; Figure S4F) and with 10μM DPI preincubation (yellow bars, N=5; Figure S4E). **(E)** Quantification of PTPRG-mCitrine catalytic cysteine oxidation in control (left) and upon knockdown of NOx complex component p22^phox^ in MCF7 cells treated with: 80ng/ml EGF-Alexa647 with or without 10μM DPI 20min preincubation, or 4mM H_2_O_2_ (mean± SEM, N=4, Figure S4F-G).

These PTPx knockdown experiments did not allow us to derive the causality between PTPx and EGFR that underlie EGFR’s response to EGF. For this, a positive perturbation is necessary which was imposed by ectopic expression of PTP_x_-mCitrine. The causality relation between PTP_x_ and EGFR was derived by analyzing the goodness of fit of the dose-response curves to the three possible modes of interaction: negative regulation, negative feedback and double negative feedback (Figure S4A, STAR Methods). A remarkable switch-like EGFR phosphorylation response was observed upon ectopic PTPRG-mCitrine expression, with a threshold of EGFR activation at around 6-7% receptor occupancy with EGF (Figure 4B top, blue line). Such a response is consistent with a double-negative EGFR-PTPRG feedback, selected by measuring the relative quality of the only three possible network motif models (Figure S4A) in representing the dose-response data (Akaike information criterion, STAR Methods). Ectopic PTPRJ-mCitrine expression flattened the dose-response curve, revealing the underlying simple negative regulation (Figure 4B bottom, blue line). Expression of the ER-bound PTPN2-mCitrine flattened the EGFR response, which could be equally well described by negative feedback or regulation (Figure 4B middle, blue line). That the knockdown of PTPRJ revealed the manifestation of the EGFR-PTPRG toggle switch indicates that the negative regulation by PTPRJ counters the switch-like EGFR phosphorylation response generated by this recursive EGFR-PTPRG interaction. The steepened dose-response upon PTPN2 knockdown (Figure 4B middle) shows a similar, although weaker role of the negative regulation by PTPN2 on the EGFR-PTPRG toggle-switch induced phosphorylation response. Consistent with siRNA-mediated knockdown, ectopic PTPRA-mCitrine expression did not affect the EGFR phosphorylation response (Figure S4D).

We investigated whether the biochemical mechanism behind the EGFR-PTPRG toggle switch originates from EGFR-induced activation of H_2_O_2_ production by NADPH-oxidases (NOX) (Bae et al., 1997), which reversibly oxidizes the catalytic cysteine in PTPs to the catalytically impaired sulfenic acid (Salmeen et al., 2003). EGFR activation by EGF increased the production of H2O2 in MCF7 cells (Figure S4B-C), so to first test whether the dose response of EGFR is affected by H_2_O_2_, we inhibited NOx activity with diphenyleneiodonium (DPI). This converted the switch-like activation observed upon PTPRG-mCitrine expression to a gradual response (Figure 4C top, yellow lines). Neither the EGF-dose response upon ectopic expression of PTPN2-mCitrine nor PTPRJ-mCitrine or PTPRA-mCitrine was affected by DPI (Figure 4C, middle and bottom and S4D). To then establish the connection between EGFR induced H_2_O_2_ production and PTPRG inhibition, we determined whether the catalytic PTPRG cysteine is oxidized upon activation of EGFR by EGF. For this, cells were incubated for 10 min with dimedone, which reacts with the sulfenylated cysteine to form a stable thioether that is detectable by an anti-dimedone antibody (Seo and Carroll, 2009). The oxidation of the catalytic cysteine (Figure S4F-G) of PTPRG increased with EGF dose (Figure 4D top, S4G), confirming the biochemical inhibitory link from EGFR to PTPRG in the toggle switch is generated by H_2_O_2_-mediated PTPRG inactivation. In contrast and consistent with the DPI experiments, neither PTPN2-mCitrine nor PTPRJ-mCitrine exhibited an EGF-dose dependent increase in catalytic cysteine oxidation (Figure 4D and S4G). To finally show that the EGF-induced oxidation of PTPRG occurs via EGFR-induced NOX activation, we knocked down the p22 subunit of NOx1-3 (p22^Phox^), resulting in a strong reduction of EGF-induced PTPRG oxidation to levels observed following DPI inhibition (Figure 4E, S4H-I).

These results therefore demonstrate that EGFR responsiveness to EGF is mainly determined by a double-negative feedback with PTPRG at the PM that is regulated by EGFR-mediated NOx-dependent production of H_2_O_2_, and modulated by PTPRJ activity at the PM and PTPN2 on the ER.

### Dynamics of the unified EGFR-PTP network

To better understand how the EGFR-PTPRG toggle switch that determines sensitivity to EGF is modulated by negative regulation by PTPRJ at the PM and negative feedback by PTPN2, we transformed the spatial scheme that describes how vesicular dynamics enables PTPs to interact with EGFR (Figure 5A) into a unified causality diagram (Figure 5B). This enabled us to explore the dynamical properties of this unified network using 3D-bifurcation analysis (Strogatz, 2000). The phosphorylation dynamics of monomeric ligandless EGFR at the PM was analyzed theoretically as function of the system’s parameters: liganded EGFR and PTPRG/EGFR expression levels.

**Figure 5.**
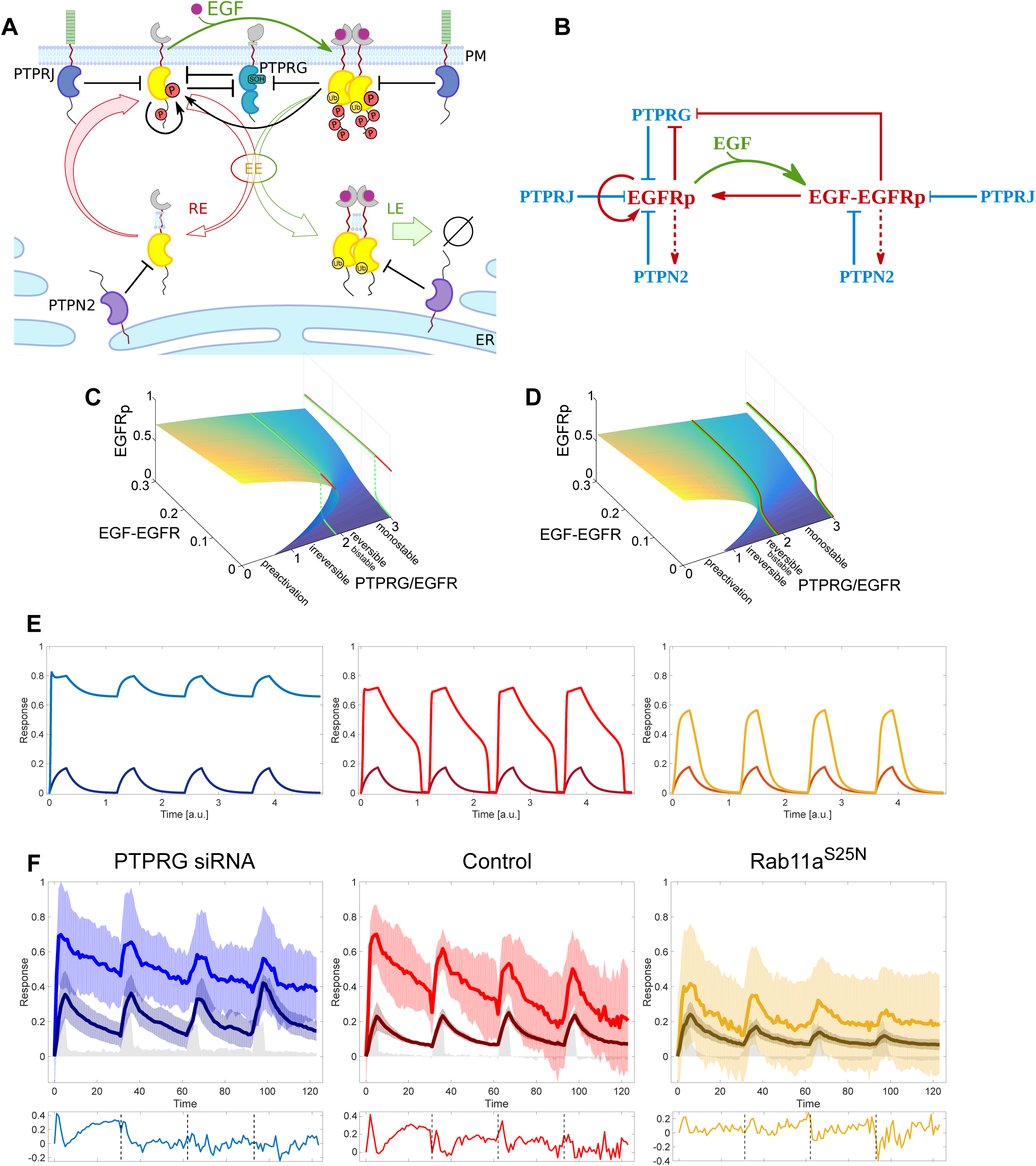
Dynamics of the spatially distributed EGFR-PTP network. **(A)** Scheme of the EGFR-PTP interaction network established through EGFR trafficking dynamics. EGFR interacts with PTPRG/PTPRJ at the PM and PTPN2 in the cytoplasm. All notations as in Figure 1K. **(B)** Causality diagram that corresponds to **(A)**. Red/blue lines: causal interactions, green arrow: ligand binding. **(C)** 3D-bifurcation diagram for double-negative EGFR-PTPRG feedback network topology at the PM, showing the dependence of monomeric EGFR phosphorylation (EGFRp) on PTPRG/EGFR expression ratio and fraction of liganded receptors. Forward (green) and backward (red) dose-response trajectories are shown for PTPRG/EGFR=1.9, with corresponding orthographic projections on the right profile plane. **(D)** 3D-bifurcation diagram as in (**C**), for the combined toggle-switch/negative regulation/negative-feedback network topology established by ligandless EGFR vesicular recycling. Projections same as in **(C)**. **(E)** Response profiles of the fractions of liganded (dark) and phosphorylated receptors (light) when the system is organized in the bistable regime (left), close to the bistability region (middle), and in the monostable regime (right), corresponding to the network topology presented in **(D)**. **(F)** Temporal traces of the fraction of ligand bound (EGF-Alexa647/EGFR-mCitrine, dark) and phosphorylated EGFR estimated by PTB-mCherry translocation to the PM (PTB-mCherry/EGFR-mCitrine, light) in live MCF7 cell expressing non-targeting siRNA (middle, n=4, N=1), following siRNA mediated knockdown of PTPRG (left, n=5, N=2), and ectopic Rab11a^S25N^ expression (right, n=16, N=2). Data was acquired at 1min-intervals following 20ng/ml 5P-EGF every 30min. Lower boxes depict the normalized differences between the fraction of phosphorylated and liganded EGFR.

We first investigated the dynamical properties of the central EGFR-PTPRG double-negative motif at the PM (Figure 5B). Together with the activation of autocatalysis on ligandless EGFR by EGF-bound EGFR, this generates bistability for a large range of PTPRG/EGFR expression. This network motif thus determines at which EGF dose EGFR collectively activates (Figure 5C, green trajectories), but impedes signal shutdown since autocatalytic EGFR activation will persist after GF removal in the bistable region (Figure 5C, red trajectories). Lowering PTPRG expression (as with knockdown) pushes the system to the pre-activated state, as demonstrated in Figure 3F. This can be alleviated by negative regulation by PTPRJ, shifting the system into the reversible bistable or even monostable region of the bifurcation diagram. Furthermore, negative regulation narrows down the bistability region and thereby shifts the EGFR phosphorylation response out of the irreversible bistability region (Figure 5D, projected trajectories). The negative feedback with PTPN2 that is established by the vesicular recycling can play a similar role to PTPRJ. However, the vesicular recycling of activated EGFR monomers that are dephosphorylated by PTPN2 in the cytoplasm is also essential to maintain sufficient EGFR at the PM for autocatalysis to manifest.

Whether and how this EGFR-PTP system will respond to time-varying cues will depend on where the system is organized in parameter space (PTPRG/EGFR). To explore how the system will respond in the different parameter regimes, we simulated EGFR responsiveness to a train of EGF pulses. If the dynamics of the EGFR-PTP system is dominated by the PTPRG-EGFR toggle switch at the PM and thus organized in the irreversible bistable region, the simulation shows that EGFR will remain “trapped” in the active state after the first EGF pulse, thereby not being able to sense subsequent EGF cues (Figure 5E, left). If the system is organized close to the bifurcation (transition to bistable region) the response dynamics exhibit biphasic behavior with a rapid decay followed by slower relaxation (Figure 5E, middle), whereas further in the monostable regime, EGFR phosphorylation closely follows the EGF input (Figure 5E, right).

To now identify where the EGFR-PTP system is poised and whether it can sense time-varying EGF signals, four subsequent 5min EGF-Alexa647 pulses followed by washout were administered every 30min to live MCF7 cells expressing EGFR-mTFP. The fraction of liganded receptor (EGF/EGFR=EGF-Alexa647/EGFR-mTFP) as well as the fraction of phosphorylated EGFR (EGFRp=PTB-mCherry/EGFR-mTFP) was ratiometrically determined at the PM as a function of time (STAR Methods). In control cells, EGFRp response relaxed in a biphasic way (with a fast and slow relaxation, light red lines Figure 5F, middle) after each EGF pulse, reminiscent of the simulated response of a system poised close to bifurcation. This differed to the relaxation of EGF/EGFR (Figure 5F middle, lower box) that approximated a more monotonic decaying function (dark red lines). The EGF/EGFR monotonic decay is due to depletion of liganded receptors from the PM by endocytosis. The more rapid activation of EGFRp with respect to EGF/EGFR at the onset of each pulse is a clear manifestation of autocatalytic EGFR amplification (Figure 5F, 1D). This shows that the EGFR-PTP system has dynamical organisation close to the bistable region, enabling both sensing, as well as robust activation upon time-varying EGF stimuli.

PTPRG knockdown results in a response to EGF pulses within the limited boundaries of the upper activated state, and never relaxes back to the basal inactivated state (light blue lines, Figure 5F left). This is consistent with the persistent/bistable EGFR phosphorylation in the absence/low level of PTPRG (Figure 3E-F). This confirms that PTPRG is a central regulator of EGFR activation dynamics through a double negative feedback motif. We also observed a subpopulation of cells (4 out of 9 cells) that relaxed back to the basal state after each EGF pulse resembling the control (Figure S4J), presumably due to variable PTPRG knockdown levels. This reflects that PTRPG concentration dictates where on the bifurcation diagram the system is organized, thereby determining the dynamics of the system. Ectopic expression of dominant negative Rab11a^S25N^ mutant impairs the vesicular recycling of EGFR monomers. This generates a lower steady state abundance of EGFR at the PM, shifting the system to the monostable regime of the bifurcation diagram by effectively increasing the system parameter PTPRG/EGFR (Figure 5D). This is apparent from the dampened phosphorylation response to a train of EGF pulses, where EGFRp follows closely the EGF/EGFR relaxation (Figure 5F right, lower box). That recycling of EGFR monomers is essential to generate a sufficient concentration of monomeric EGFR for autocatalytic amplification of phosphorylation after each EGF pulse is apparent from the strong decrease in both autocatalytic EGFR activation (Figure 5F right), as well as the dampening of both EGFRp and EGF/EGFR after each pulse. In this case, the system loses its robustness in response to time varying stimuli and becomes more rapidly insensitive to upcoming EGF pulses (Figure 5F, light orange lines). How long the system can respond to time-varying EGF stimuli generally depends on the total amount of expressed EGFR that is recycling in the cell, and how quickly this pool is depleted by the unidirectional trafficking of liganded EGFR, determined by the magnitude of EGF stimuli.

## Discussion

By relating EGFR-mTFP phosphorylation at physiological expression levels to systematic opposed genetic perturbations by PTP knockdown and ectopic expression, we not only identified which PTPs regulate EGFR phosphorylation, but also derived a measure for the specific reactivity of these PTPs towards EGFR. This enabled the identification of the juxtaposed receptor-like PTPRG/J at the PM and ER-bound PTPN2 (TCPTP) as major PTPs that regulate EGFR phosphorylation dynamics. The additional non-redundant PTPs with lower specific activity (PTPRA, DUSP3) that were identified in the CA-FLIM screen point at additional modulatory layers that control EGFR phosphorylation. Here, we could show that PTPRA helps to suppress autonomous activation of recycling EGFR at the PM late after stimulus and that the soluble DUSP3 that consist solely of a phosphatase domain can dampen overall EGFR phosphorylation by the regulation of its expression (Andersen et al., 2001).

In order to understand how the PTPs on juxtaposed membranes regulate sensitivity to growth factors and subsequent EGFR phosphorylation response, it was important to distinguish the vesicular and phosphorylation dynamics of ligandless and liganded EGFR (Figure 1F-J). The receptor that binds ligand is dimerized and ubiquitinated by cCbl binding, providing the signal for unidirectional traffic to the LE (Baumdick et al., 2015) to be degraded in lysosomes. During vesicular trafficking, liganded EGFR moves along an increasing PTPN2 activity that is imposed by the increasing amount of ER membranes towards the perinuclear area. The high phosphatase activity that is necessary to dephosphorylate kinase active EGFR dimers is the reason why most of PTPN2 activity is spatially segregated from the PM where it would otherwise strongly suppress EGFR activation upon ligand exposure. In contrast, the relative mRNA expression of PTPRG and PTPRJ with respect to PTPN2 (PTPR/ER-PTP mRNA~0.045, Figure S1A) show that the high specific activity of PTPRs at the PM is compensated for by lower endogenous expression to avoid complete suppression of EGFR phosphorylation upon EGF stimulation. Since EGF binding promotes endocytosis of the receptor, the interaction between EGFR and PTPN2 (Figure 5A) becomes conditional on EGFR activity, effectively generating a negative feedback. In this case, EGFR signal duration is determined by vesicular trafficking rates and occurs within tens-of-minutes. This long time scale cannot be achieved by a rapid, diffusion controlled biochemical reaction, such as recruitment of cytosolic SHP1/2 to phosphorylated EGFR (Grecco et al., 2011). The major function of PTPN2 is therefore tightly linked to vesicular trafficking of liganded EGFR to control the signal duration of ligand activated receptors. The only other ER-bound PTP is PTPN1 (Frangioni et al., 1992), which manifested its EGFR dephosphorylating activity only upon ectopic expression in our CA-FLIM screen. This shows that PTPN1 is redundant with respect to PTPN2, and likely performs a similar functionality in regulating EGFR’s temporal response (Baumdick et al., 2015; Haj et al., 2002; Yudushkin et al., 2007)

In order to understand the role of PTPRs at the PM, it was necessary to consider the autocatalytic activation of ligandless EGFR. The manifestation of EGFR autocatalysis was previously reported (Reynolds et al., 2003; Tischer and Bastiaens, 2003), but the more recently described regulatory Y_845_ in the kinase activation loop, which stabilizes the active conformation of the receptor upon phosphorylation (Shan et al., 2012), could provide the basis for this mechanism. Autocatalysis is thereby established by direct autophosphorylation or indirect phosphorylation by Src (Sato et al., 1995) that is in turn activated by EGFR (Osherov and Levitzki, 1994), effectively generating the same positive feedback. The single-cell dose response experiments demonstrated that ligand-bound EGFR can trigger autocatalytic phosphorylation on ligandless receptors, thereby establishing an activatory link from ligand-bound to ligandless EGFR (Figure 1D). These monomeric species constitutively recycle through the cytoplasm where they are dephosphorylated by PTPN2 before repopulating the PM. Recycling of EGFR monomers is therefore essential to generate a sufficient concentration of receptors for autocatalytic phosphorylation amplification at the PM. We could show that the tumor suppressor PTPRG (Kwok et al., 2015) preferentially dephosphorylates EGFR monomers and that its enzymatic activity is coupled to that of phosphorylated EGFR via NOX-dependent H_2_O_2_ production. This EGF-dependent inactivation of PTPRG activity through the oxidation of its catalytic cysteine thereby establishes a double negative feedback at the PM that generates switch-like EGFR activation at a threshold EGF concentration. This system has the property to be robust to noise in extracellular GF concentrations but also has the fragile property of being irreversible after activation. This was reflected in EGFR preactivation upon PTPRG knockdown and is in line with PTPRG being a tumor suppressor. The negative regulation of EGFR by PTPRJ opposes the tendency of the EGFR-PTPRG toggle switch to induce irreversible EGFR phosphorylation by pushing the system out of bistability.

PTPN2 on the ER also negatively modulates the EGFR phosphorylation dynamics generated by the EGFR-PTPRG toggle switch, both through a small fraction of its total enzymatic activity at ER-PM contact sites (Lorenzen et al., 1995) as well as by dephosphorylating recycling EGFR monomers in the cytoplasm. This establishes a dynamically coupled EGFR-PTPN2 negative feedback (Figure 5A,B).

Growth factor receptors are the ‘sensory organs’ of cells that perceive time varying growth factor stimuli that can lead to a variety of cellular responses. Here we have shown that an EGFR-PTPRG toggle switch at the PM is coupled to a negative regulation by PTPRJ at the PM and a cytosolic EGFR-PTPN2 negative feedback (Figure 5B). The dynamical organization of the system is poised such that EGFR is only activated when physiological EGF concentrations are present in the extracellular medium and generates a system that can robustly respond to time-varying stimuli in a non-stationary environment. Within this process, signaling has to be terminated after the transduction takes place, while continuously repopulating the plasma membrane with receptors that can sense upcoming cues. Given the role of vesicular dynamics in the regulation of EGFR activation and signaling, changes in its vesicular dynamics may represent a mechanism through which environmental inputs such as cell-cell contact can influence the cellular response to EGF stimulation, generating contextual plasticity in growth factor signaling.

## Author contributions

P.I.H.B. conceived the project. P.I.H.B. and A.K. developed the dynamical system concepts and supervised the study. H.E.G. designed and S.F., P.R. –N., H.E.G., K.C.S. and A.M. performed the FLIM screening experiments, A.M., A.S., K.C.S., W.S. and M.B – EGFR phosphorylation and trafficking experiments, M.B. – anisotropy experiments, A.S., R.S. and Y.B. – single-cell dose-response experiments, M.S.J., R.S. and A.M. – oxidation assays, A.S. and W.S. performed the EGF-pulse experiments and L.R. and J.L. generated the PTP cDNA plasmid library. A.S., H.E.G., K.C.S. and A.K. performed image- and data-analysis. P.I.H.B., A.K. and A.S. developed the theoretical models. P.I.H.B. and A.K. wrote the manuscript with help of A.S. and A.M.

## Acknowledgments

The project was partly funded by the European Research Council (ERC AdG 322637) and the European Commission (Grant 278568) to P.I.H.B. P.R.N was supported by Marie Curie Actions (IEF 219743). The authors thank Kirsten Michel for help with biochemical experiments, Dr. Jan Hübinger for help with setting up the EGF-pulse instrumentation and Drs. Astrid Krämer, Peter Bieling and Malte Schmick for critically reading this manuscript.

## Supplemental figure legends

**Figure S1.**
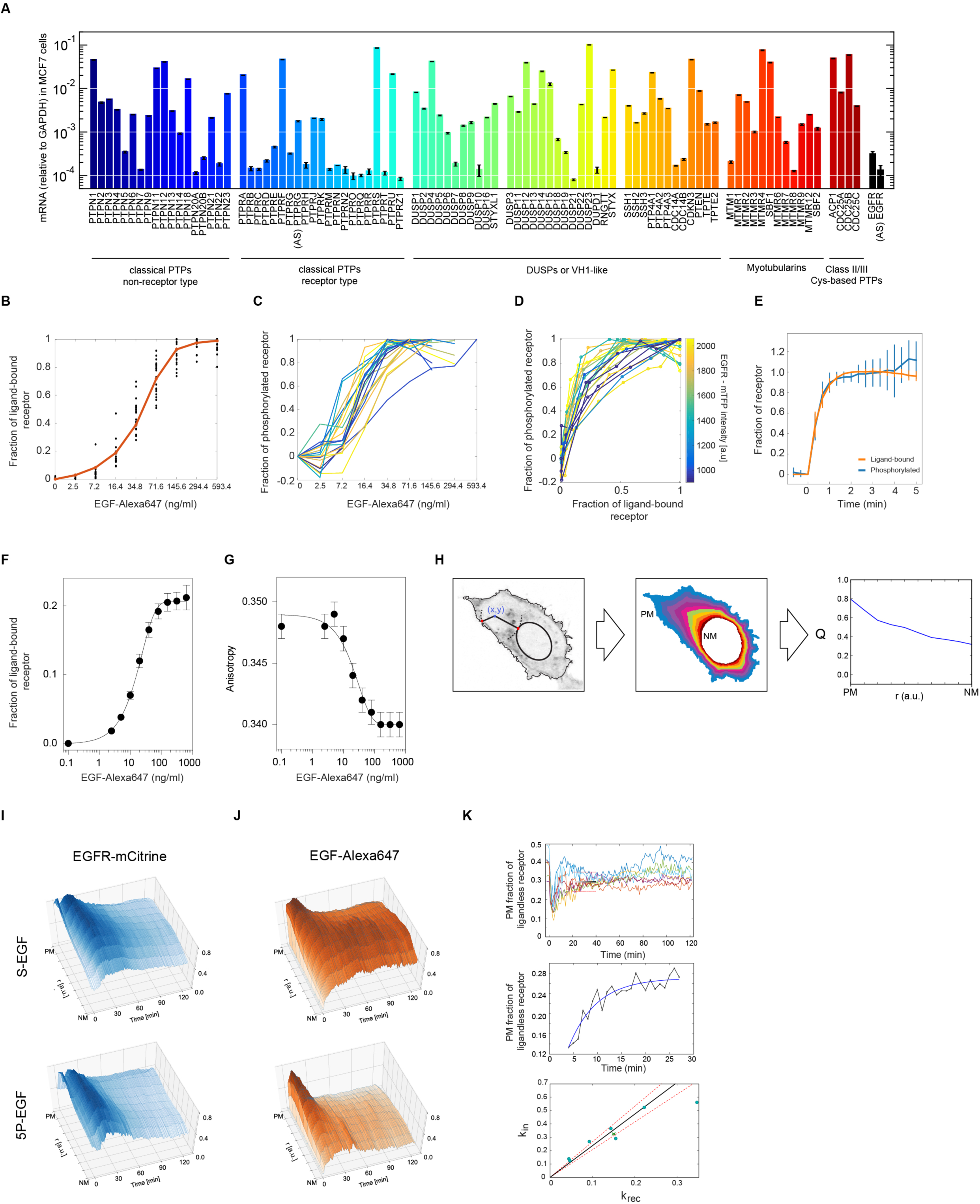
EGFR phosphorylation and trafficking dynamics, related to Figure 1. **(A)** PTP_X_ and EGFR mRNA expression in MCF7 cells measured by microarray analysis (relative to GAPDH mRNA). AS: anti-sense. **(B)** Estimating EGFR-mTFP occupancy with EGF-Alexa647 in live cells: EGF-Alexa647/EGFR-mTFP quantified in single cells (data points) upon increasing EGF-Alexa647 doses was fitted (line) with a receptor binding kinetics model (STAR Methods). **(C)** Fraction of phosphorylated receptor, quantified by PTB-mCherry translocation to the PM (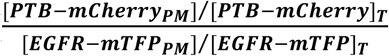 star Methods) for different EGF-Alexa647 doses (single cell profiles). **(D)** Single cell EGFR-mTFP phosphorylation dose response profiles (average shown in Figure 1D). The estimated fractions of phosphorylated vs. liganded EGFR-mTFP are plotted and color-coded by the average EGFR-mTFP fluorescence intensity per cell. **(E)** Quantification of PTB-mCherry translocation kinetics to PM-localized EGFR-mTFP. MCF7 cells were stimulated with a saturating EGF-Alexa647 dose (320ng/ml) and successive images were acquired every 20s (n=10). Translocated PM fraction of PTB-mCherry (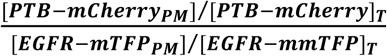) converged to a steady state level in ~1.5min, which was within the time frame of successive EGF-Alexa647 dose administration (Figure 1C-D). **(F)** Estimation of EGFR-QG-mCitrine occupancy with EGF-Alexa647 in live cells (n=30, N=3) from fluorescence anisotropy microscopy is equivalent to the corresponding estimation from confocal microscopy with EGFR-mTFP (Figure S1B). **(G)** Dependency of the EGFR dimerization state with increasing EGF-Alexa647 doses determined by fluorescence anisotropy microscopy (n=30, N=3). **(H)** Dimensionality reduction from Cartesian (x, y) to normalized radial (r) distribution of quantity (Q) between the plasma (PM) and the nuclear (NM) membrane. **(I)** Average spatial-temporal maps (STMs) of EGFR-mCitrine intensity obtained from live cells stimulated with 200ng/ml S-EGF (n=16, N=3; top) and 5P-EGF (n=14, N=2; bottom). **(J)** Corresponding STMs of EGF-Alexa647 fluorescence. **(K)** Quantification of recycling dynamics of ligandless EGFR-mCitrine upon 200ng/ml 5P-EGF. Top: PM fraction of ligandless EGFR-mCitrine in single cells. Model-based estimation of the steady state level (95% confidence bounds; see STAR Methods) is shown with black line (inside red dashed lines). Middle: compartment-model-based fitting on 4-28min interval for the cells shown in the bottom panel of Figure 1G (estimated rates: k_in_=0.125, k_rec_=0.046, STAR Methods). Bottom: Linear dependency between (k_in_, k_rec_) reflects that similar PM steady state levels of ligandless EGFR are maintained by recycling in single cells (x: average (k_in_, k_rec_); STAR Methods).

**Figure S2.**
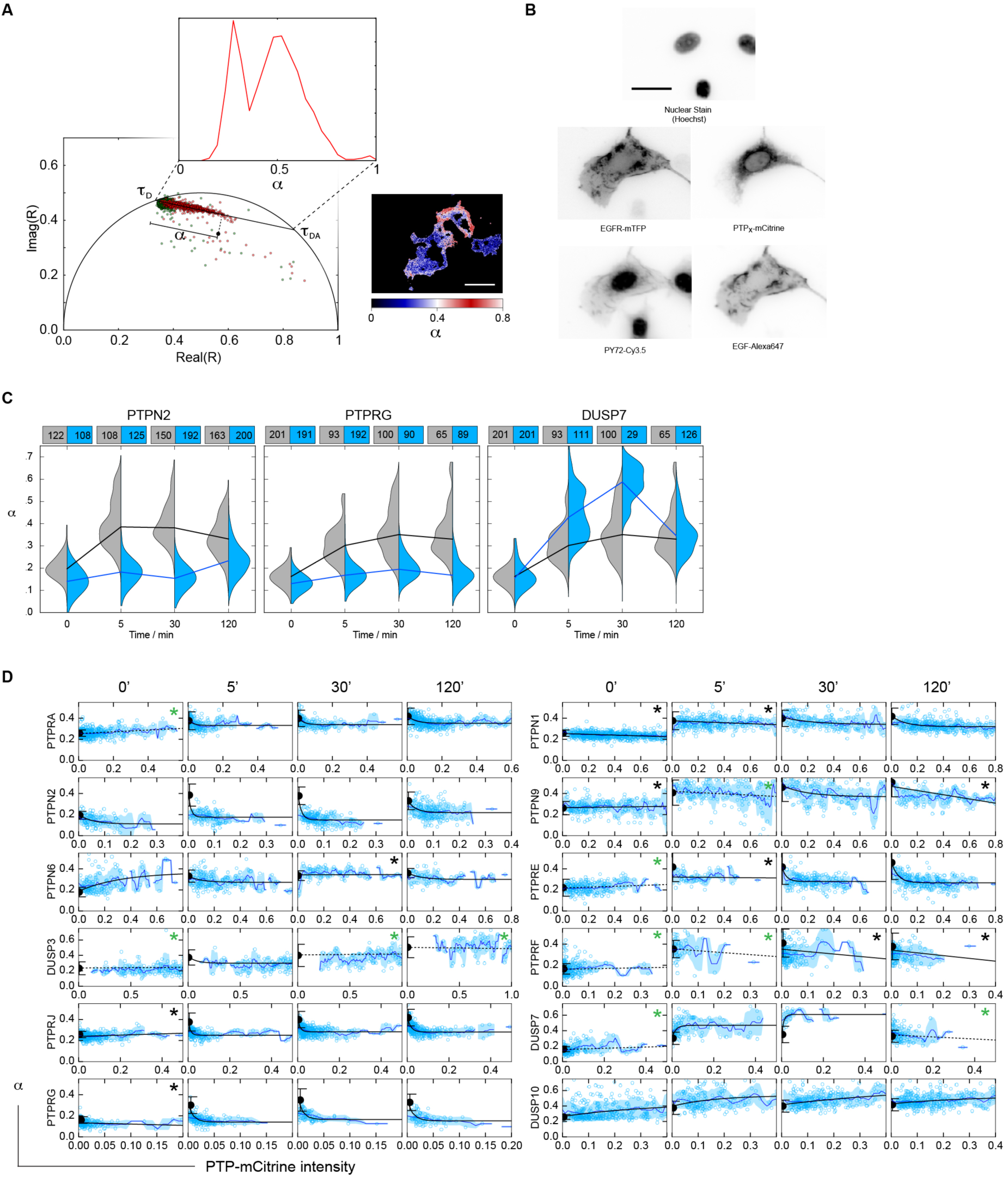
CA-FLIM and related quantities, Related to Figure 2. **(A)** Linear fit of the fluorescence emission Fourier coefficients (R) in the complex plane yielding the global EGFR-mTFP lifetimes in presence (τ_DA_) and absence (τ_D_) of FRET. Fraction of phosphorylated EGFR-mTFP bound by PY72-Cy3.5 (α) in each pixel was calculated from the projection onto the τ_D_-τ_DA_ line segment. An exemplary α-histogram (inset) and a spatially resolved α-map (right) are shown. Scale bar: 20μm. **(B)** Representative images obtained in CA-FLIM screen: Hoechst (nuclear stain), EGFR-mTFP, PTP_X_-mCitrine, PY72-Cy3.5 and EGF-Alexa647 fluorescence. Scale bar: 10μm. **(C)** Exemplary temporal EGFR-mTFP phosphorylation profiles (grey, control) upon ectopic PTP_X_-mCitrine expression (PTP_X_=PTPN2, PTPRG, DUSP7; blue). The violin plots show the α distributions from single cells stimulated with 200ng/ml 5P-EGF (number of cells denoted on top of the plots, medians at different time points are connected by a black line). **(D)** α vs. PTP_X_-mCitrine single cell fluorescence scatter plots. Black circle: mean α_ctr_±SD; black lines: exponential fits (* - linear fit for weak dependence, green asterisk: distributions of α_ctr_ and α_PTP_ did not significantly differ); blue lines with error bounds: moving averages with standard deviations. The slope of the exponential (or linear) fit at the intercept is defined as the relative specific PTP_X_-mCitrine activity in Figure 2C (middle).

**Figure S3.**
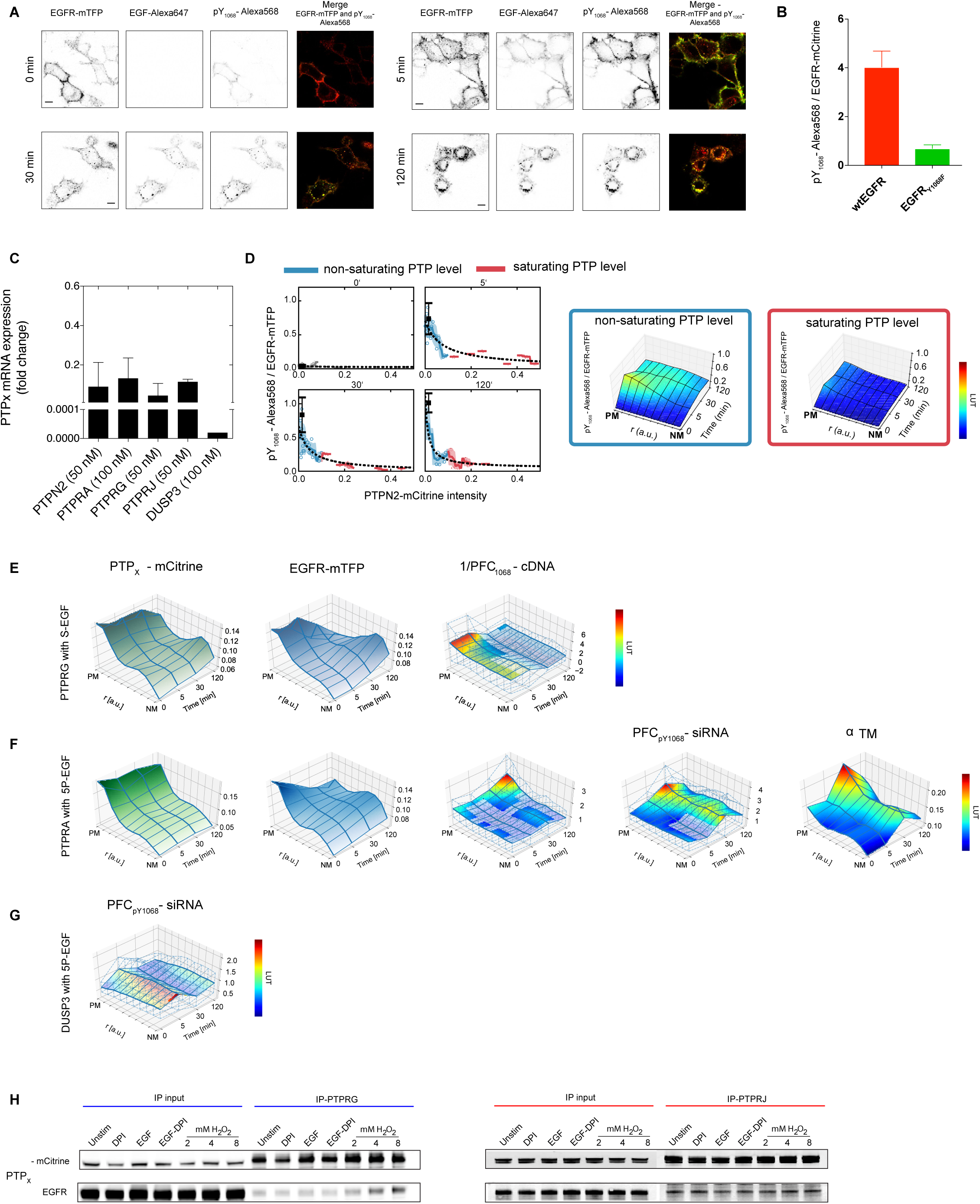
PTPs shape EGFR phosphorylation patterns, Related to Figure 3. **(A)** Representative images of EGFR-mTFP, EGF-Alexa647, anti-pY_1068_-Alexa568 fluorescence and overlay of pY_1068_ (yellow) and EGFR-mTFP (red) prior to and after 5, 30 and 120min of 5P-EGF stimulation (200ng/ml) of MCF7 cells. Scale bar: 50μm. **(B)** Binding of pY_1068_-Alexa568 antibody to Y_1068_ on EGFR-mTFP and EGFR^Y1068F^-mTFP reflects its specificity for the corresponding tyrosine phosphorylation site (mean±SD, n~100, N=1). **(C)** mRNA expression fold change of PTPN2, PTPRG, PTPRJ, PTPRA and DUSP3 in MCF7 cells after 24h transfection with 50nM respective siRNA. The values are relative to mRNA levels of the respective gene in cells treated with 50nM nontargeting siRNA for 24hr (N=2, and for DUSP3 N=1). **(D)** Left: Example of pY_1068_-Alexa568/EGFR-mTFP vs. PTPN2-mCitrine fluorescence intensity scatter plots used to determine the PTPN2-mCitrine fluorescence intensity threshold at which saturation of EGFR dephosphorylation occurs. Blue/red circles represent single cells below and above the PTPN2-mCitrine fluorescence intensity threshold, respectively; solid lines and shaded bounds: corresponding moving averages and standard deviation. The data was fitted with a function depicting the dependency of pY_1068_/EGFR-mTFP on PTP_X_-mCitrine intensity (steady state EGFRp assumption, STAR Methods). Right: STMs of pY_1068_/EGFR-mTFP averaged from cells below (blue box) and above (red box) PTPN2-mCitrine fluorescence intensity threshold. LUT: look-up table. **(E)** Effect of PTPRG-mCitrine expression (left) on STMs of EGFR-mTFP fluorescence (middle) and phosphorylation fold-change (1/PFC_pY1068_-cDNA, right) reflecting the relative PTPRG-mCitrine reactivity towards pY_1068_ for cells stimulated with 200ng/ml S-EGF (n~30). LUT: look-up table. (F) Columns 1-3: effect of PTPRA-mCitrine expression (Column 1) on STMs of EGFR-mTFP localization (Column 2) and phosphorylation fold-change (1/PFC_pY1068_-cDNA, Column 3), which reflects the relative PTPRA-mCitrine reactivity towards pY_1068_ (n~60, N=3). Column 4: effect of siRNA-mediated PTPRA knockdown on EGFR-mTFP phosphorylation fold change (PFC_pY1068_-siRNA, n~45, N=3). Column 5: STM of fraction of EGFR-mTFP interacting with PTPRA ^C442S^-mCitrine trapping mutant (α_™_, n=15-30). LUT: look-up table. **(G)** Effect of siRNA-mediated DUSP3 knockdown on EGFR-mTFP phosphorylation fold change (PFC_pY1068_-siRNA, n~40, N=3). In (F-G), cells were stimulated with 200ng/ml 5P-EGF; transparent areas in (E-G): non-significant PFCs, p >0.05. LUT: look-up table. **(H)** Identifying and characterizing the interaction between EGFR and PTPRG/J-mCitrine by co-immunoprecipitation. Co-immunoprecipitated EGFR (second row) following PTPRG- (left) or PTPRJ-mCitrine (right) pull down from MCF7 cells prior to and after treatment with DPI (20min, 10μM), EGF-Alexa647 (10 min, 200ng/ml), DPI pretreatment (20min, 10μM) followed by EGF-Alexa647 (10 min, 200ng/ml) and H_2_O_2_ (10 min; 2, 4 or 8 mM) by western blotting. IP input: total expressed protein, IP-PTPRG/J: PTPRG/J-mCitrine immunoprecipitated by anti-GFP antibody.

**Figure S4.**
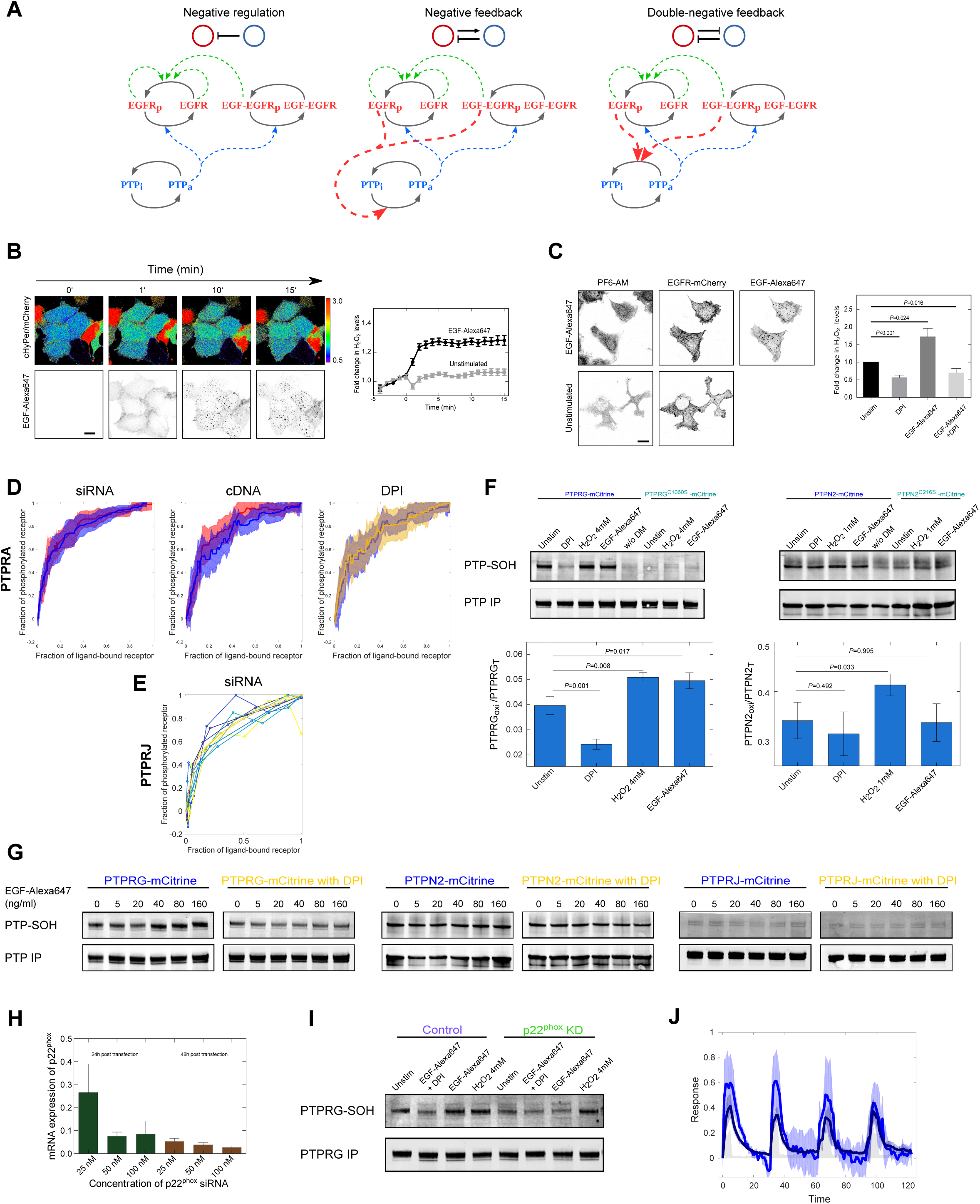
Quantifying EGF-induced H_2_O_2_ production and ROS mediated oxidation of catalytic cysteines in PTPs, Related to Figure 4. **(A)** Possible EGFR-PTP network motifs. Solid arrows: molecular state transitions (p: phosphorylation, i: inactive, a: active), dashed arrows: causal links. Left: negative regulation, middle: negative feedback, right: double negative feedback. Top row: schematic model representations. **(B)** Left: Representative pseudo-coloured image series of cHyPer3/mCherry fluorescence ratio (upper row), normalized to that at 0min in each MCF7 cell. Images were acquired every minute for 15min after 320ng/ml EGF-Alexa647 stimulation (lower row). Scale bar: 20μm. Right: Corresponding quantification of relative H_2_O_2_ levels (cHyPer3/mCherry fluorescence ratio ± SEM) upon EGF-Alexa647 (black line, n=11 cells) or vehicle (unstimulated) administration (grey line, n=9 cells). **(C)** Left: fluorescence images of the H_2_O_2_-sensitive probe PF6-AM (left), EGFR-mCherry (middle) and EGF-Alexa647 (right) in MCF7 cells with (upper row) and without (lower row) 320ng/ml EGF-Alexa647 stimulation. Scale bar: 20μm. Left: corresponding quantification of H_2_O_2_ fold-change (mean PF6-AM fluorescence ± SEM from 10 fields of view) upon administration of DPI (10 min, 10μM), EGF-Alexa647 (10 min, 320 ng/ml) or a combination of both. **(D)** Dose-response of EGFR-mTFP phosphorylation (control, red) is not affected upon siRNA-mediated PTPRA knockdown (blue, p=0.823, n=14, N=3, left) or PTPRA-mCitrine co-expression (blue, p=0.225, n=34, N=5, middle). NOx-inhibition by DPI (10μM, 30min pre-incubation) has no effect (yellow, p=0.937, n=38, N=4, right) on the dose-response of EGFR phosphorylation upon ectopic PTPRA-mCitrine expression (blue, same as in middle plot). **(E)** Representative dose response curves of EGFR-mTFP phosphorylation in single cells upon siRNA-mediated PTPRJ knockdown. The corresponding average is shown in Figure 4A, bottom. **(F)** Quantification of cysteine oxidation (PTP-SOH) in PTPRG-mCitrine (left) or PTPN2-mCitrine (right) to cysteine sulfenic acid by dimedone conjugation, detected by anti-sulfenic acid modified cysteine antibody. Left: detection of oxidized cysteines in immunoprecipitated PTPRG-mCitrine from MCF7 cells treated for 10 min with DPI (10μM), EGF-Alexa647 (80ng/ml) or 4mM H_2_O_2_ by western blotting. The weak PTP-SOH bands in the corresponding lanes of the PTPRG^C1060S^-mCitrine mutant show that it is the catalytic cysteine is oxidized. Right: Oxidation of cysteines in PTPN2-mCitrine and PTPN2^C216S^-mCitrine detected using same conditions as above (with the exception of the H_2_O_2_ dose). **(G)** Cysteine oxidation of PTPRG-mCitrine (left), PTPN2-mCitrine (middle) and PTPRJ-mCitrine (right) upon 10min stimulation with increasing EGF-Alexa647 doses (0-160ng/ml) with and without pretreatment (20 min, 10μM) (representative WBs). **(H)** mRNA expression of p22^phox^ determined by RT-PCR after 24h or 48h of siRNA transfection with different concentrations (N=2). (I) Representative WB of cysteine oxidation in PTPRG-mCitrine upon non-targeting (control) or p22^phox^ siRNA transfection. Quantification given in Figure 4F. (J) Temporal traces of the fraction of ligand bound (EGF-Alexa647/EGFR-mCitrine, dark) and phosphorylated EGFR estimated by PTB-mCherry translocation to the PM (PTB-mCherry/EGFR-mCitrine, light) in live MCF7 cell following siRNA mediated knockdown of PTPRG (n=4, N=2). Data was acquired at 1min-intervals following 20ng/ml 5P-EGF every 30min.

## STAR Methods

### Contact for Reagent and Resource Sharing

Further information and requests for resources and reagents should be directed to and will be fulfilled by the Lead Contact, Prof. Dr. Philippe I. H. Bastiaens

### Experimental Model and Subject Details

#### Cell culture

MCF-7 cells (ECACC, Cat. No. 86012803) and MCF7 cells stably expressing EGFR-EGFP (EGFR^-EGFP^ MCF7) were cultured in Dulbecco’s modified Eagle’s medium (DMEM) (PAN Biotech), supplemented with 10% heat-inactivated fetal bovine serum (FBS) (PAN Biotech), 10mM glutamine (PAN Biotech) and 1% Non-Essential Amino Acids (PAN Biotech) at 37°C with 5% CO_2_. MCF10A (ATCC-CRL 10317) were grown in DMEM/F12 media supplemented with 5% horse serum, 20ng/ml EGF (Sigma-Aldrich), 0.5μg/ml hydrocortisone (Sigma #H-0888), 100ng/ml cholera toxin (Sigma), 10μg/ml insulin (Sigma) and 1% glutamine. MCF7 and MCF10A cells were authenticated by Short Tandem Repeat (STR) analysis (Leibniz-Institut DSMZ). Cells were regularly tested for mycoplasma contamination using MycoAlert Mycoplasma detection kit (Lonza).

### Method Details

#### Expression plasmid library

The p2297-OPIN(n)mCitrine (Berrow et al., 2007) and p2150-OPIN(c)mCitrine (Berrow et al., 2007) vectors without His6-Tag were used to generate N-or C-terminally tagged PTP_X_–mCitrine expression constructs. See Table S2 for PTP_X_ constructs with mRNA reference ID, source of the cDNA/ORF, vector, sequence of the Ligation-Independent-Cloning-(LIC) primers and any sequence discrepancies. To obtain PTP ORFs from human cell lines, mRNA was isolated with the RNeasy Maxi and Oligotex mRNA Midi Kit (QIAGEN) followed by cDNA synthesis using the AffinityScript Multiple Temperature cDNA Synthesis Kit (Agilent). The cloning of ORF into the pOPIN vector was done with a combination of ‘in vivo cloning’ (Oliner et al., 1993) and “sequence and ligase independent cloning (SLIC)” (Li and Elledge, 2007) by the Dortmund Protein Facility. The PCR reaction comprised of LIC primers and Phusion Flash High-Fidelity PCR Master Mix (Thermo Fisher Scientific) or Herculase II Fusion DNA Polymerase (Agilent). PTP_x_-pOPIN sequences were validated using BigDye^®^ Terminator v3.1 Cycle Sequencing Kit (Thermo Scientific). Plasmids were extracted from transformed E.coli XL - 10 Gold ultracompetent cells using a high content PureYield plasmid Midiprep System (Promega) and NucleoBond^®^ Xtra Midi EF (Macherey-Nagel). Trapping mutants were generated for PTPs listed in Fig.2b. See Table S2 for site of mutation and the respective LIC and mutagenesis primer pairs Table S2. Mutations were introduced into the WT PTP_X_ cDNA by an overlap extension PCR and later cloned into the respective vector using LIC. EGFR-mTFP-N1, was generated from EGFR-mCitrine-N1 (Baumdick et al., 2015) using AgeI and NotI restriction enzymes to exchange mCitrine with mTFP1. EGFR^Y845F^-mCitrine was generated by site-directed mutagenesis with the primers listed in Table S2. The EGFR-QG-mCitrine construct has been previously described (Baumdick et al., 2015). The constructs of PTB-mCherry, EGFR-mCherry and cCbl-BFP were described previously (Fueller et al., 2015). cHyPer3 (Bilan et al., 2013) plasmid was kindly provided by Prof. Vsevolod Belousov, Shemyakin–Ovchinnikov Institute of Bioorganic Chemistry, Moscow.

#### Antibodies

Primary antibodies: Mouse monoclonal antibody PY72 (Glenney et al., 1988) (InVivo Biotech Services, Henningsdorf, Germany), rabbit anti EGFR pY_1068_ (Cell Signaling; 1:400), goat anti EGFR (R&D Systems; 1:300). Secondary antibodies: Alexa Fluor^®^ 568 donkey anti-rabbit IgG (Life Technologies, 1:200), Alexa Fluor^®^ 568 donkey anti-mouse IgG (Life Technologies, 1:200), Alexa Fluor^®^ 647 donkey anti-goat IgG (Life Technologies, 1:200), Alexa Fluor^®^ 647 chicken anti-mouse IgG (Life Technologies, 1:200), Alexa Fluor^®^ 647 donkey anti-rabbit IgG (Life Technologies, 1:200), IRDye^®^ 800CW Donkey anti-Rabbit IgG (Licor, 1:10000), IRDye^®^ 680RD Donkey anti-Mouse IgG (Licor, 1:10000).

#### hEGF-Alexa647

The His-CBD-Intein-(Cys)-hEGF-(Cys) plasmid (Sonntag et al., 2014) was kindly provided by Prof. Luc Brunsveld, University of Technology, Eindhoven. Human EGF was purified from E. coli BL21 (DE3) and N-terminally labeled with Alexa647-maleimide as described previously (Sonntag et al., 2014) and stored in PBS at −20°C.

#### PY72-Cy3.5 labelling

Cy3.5^®^ NHS ester (GE Healthcare) was dissolved in 10μl of dried N,N dimethylformamide (SERVA Electrophoresis). For each reaction, 15μl of 1M Bicine (pH 9.0) and a 10-fold molar excess (to PY72) of Cy3.5 were added to 100μl PY72 (0.25mg/ml) in PBS. After 20min in the dark reaction was terminated by adding 6μl of 0.2M Tris buffer (pH 6.8). Free dye was removed by using 7K Zeba Spin Desalting Columns (Thermo Scientific). The absorption (A) of the filtrate was measured at 280nm (PY72) and 581nm (Cy3.5). For immunostaining, labelled antibody (30μg/ml in PBS) with dye to protein ratio of 3 - 5 was used. 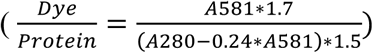

#### Transfection and EGF treatment

3×10^4^ MCF7 cells were seeded per well in an 8-well Lab-Tek chamber (Nunc). After 7-8h of seeding, cells were transfected with 0.125μg of each plasmid (EGFR-mTFP, PTP_X_-mCitrine or cCBL-BFP) using FUGENE6 (Roche Diagnostics) and incubated overnight. Before EGF stimulation, cells were serum starved with supplemented DMEM (see above) for 6h. The cells were stimulated with a sustained or a 5min-pulse of 200ng/ml EGF-Alexa647. Cells were chemically fixed with Roti^®^ Histofix 4% (Carl Roth) for 20min, washed three times with PBS and then permeabilized with 0.1% Triton-X/PBS (SERVA Electrophoresis) for 15min. Cells were stored in PBS at 4°C before immunostaining. For live cell EGFR trafficking experiments, MCF7 cells were seeded at ~2×10^4^ cells/well in an 8-well Lab-Tek chamber (S-EGF, 200ng/ml) or ~10^5^ cells/well in a 6-well dish with a cover slide (Masip et al., 2016) (5P-EGF, 200ng/ml) and transfected after 24h with a total of 0.15-0.22μg (8-well) or 1-2.5μg (6-well) of EGFR-mCitrine, PTB-mCherry and cCbl-BFP expression plasmids. In experiments requiring siRNA transfection, cells were transfected 6h before cDNA transfection with DharmaFECT1 (Dharmacon) according to the manufacturer’s instructions. Before EGF stimulation, cells were serum starved with supplemented DMEM for at least 6h. For Live cell dose response experiments, MCF7 cells were seeded at ~2×10^4^ cells/well in an 8-well Lab-Tek chamber and transfected after 24h using FUGENE6 (Roche Diagnostics) with 0.15μg TagBFP, EGFR-mTFP/EGFR-mCitrine/EGFR^Y845F^-mCitrine, PTB-mCherry and PTPRG/PTPN2-mCitrine (where applicable) per well. Before EGF stimulation, cells were serum starved with supplemented DMEM with 0.5% FCS for at least 6h. For live cell dose response anisotropy experiments, 1.5×10^5^ MCF7 cells were seeded in a MatTek (MatTek Corporation) dish and transfected with 1.5μg EGFR-QG-mCitrine using FUGENE6 (Roche Diagnostics) after 24h. For the time-lapse anisotropy experiment with 5P-EGF or S-EGF stimulation, 1.5×10^5^ MCF7 cells were seeded in a MatTek (MatTek Corporation) dish and transfected after 24h with 1.6μg EGFR-QG-mCitrine and 1μg cCbl-BFP using FUGENE6 (Roche Diagnostics). Before EGF stimulation, cells were serum starved with supplemented DMEM with 0.5% FCS for 6h.

#### Reverse transfection for CA-FLIM

siRNA and cDNA arrays were prepared and stored according as described previously (Grecco et al., 2010). Each array constituted of 384 siRNA or cDNA reverse transfection spots printed on a NaOH treated glass slide of 1-well Lab-Tek chamber (Nunc). Along with other components (Grecco et al., 2010), the transfection-spotting mixture comprised of either 0.67μM siRNA Smart-Pools (Dharmacon, Table S3) for the siRNA array or 0.5μg of EGFR-mTFP and PTP_X_-mCitrine plasmid for the cDNA array. For siRNA arrays 2.5×10^5 EGFR-EGFP^ MCF7 cells and for cDNA arrays 3×10^5^ MCF7 cells were seeded and incubated for 48h. Before EGF stimulation, cells were serum starved with supplemented DMEM (see above) without FCS for 6h. The cells were stimulated for 5, 30 or 120min with a sustained or 5min-pulse of 200ng/ml EGF-Alexa647. Cells were fixed chemically with Roti^®^ Histofix 4% (Carl Roth) for 20min, washed three times with PBS and then permeabilized with 0.1% Triton-X/PBS (SERVA Electrophoresis) for 15min. Cells were stored in PBS at 4°C before immunostaining.

#### Identifying the optimal siRNA concentration

2×10^5^ of MCF7 cells were seeded in each well of a 6-well tissue culture dish and transfected after 24h using 50nM siRNA specific for PTPN2, PTPRG, PTPRJ, CYBA or non-targeting control siRNA with Dharmafect1 according to the manufacturer’s instructions. RNA was isolated 24h after transfection using the Quick-RNA MicroPrep kit (Zymo Research, Freiburg, Germany). For quantification of mRNA expression levels of interest, 1μg input RNA was used for reverse transcription using the High Capacity Reverse Transcription kit (Applied Biosystems) according to the manufacturer instructions. Commercially available TaqMan assays (Thermo Fisher), PTPN2(Hs00959888_g1), PTPRG(Hs00892788_m1), PTPRJ(Hs01119326_m1), GAPDH(Hs02786624_g1), CYBA(Hs00609145_m1) were used to detect the amplicons after each cycle of a qPCR reaction ran in an IQ5 real-time PCR system cycler (Bio-Rad). Cycling condition were as follows: 40 cycles of 95°C for 10s and 57°C for 30s. Data were analysed using the ΔΔCt method for determination of relative gene expression by normalisation to an internal control gene (GAPDH), and fold expression change was determined compared to the control siRNA sample.

#### In-cell westerns

MCF7 and MCF10A cells were seeded on black, transparent bottomed 96-well plates (3340, Corning, Hagen, Germany) coated with poly-L-lysine (P6282, Sigma Aldrich), transfected 24h later when required and starved for 18h in DMEM containing 0.5% FCS prior to stimulation. After stimulation, cells were fixed with Roti-Histofix 4% (Carl Roth, Karlsruhe, Germany) for 5min at 37°C and permeabilized with 0.1% Triton X-100 (v/v) for 5min at room temperature. Samples were incubated in Odyssey TBS blocking buffer (LI-COR Biosciences, Lincoln, NE, USA) for 30min at room temperature. Primary antibodies were incubated overnight at 4°C and secondary antibodies (IRDyes, LI-COR Biosciences) were incubated in the dark for 1h at room temperature. All wash steps were performed with TBS (pH 7.4). Intensity measurements were made using the Odyssey Infrared Imaging System (LI-COR Biosciences). Quantification of the integrated intensity in each well was performed using the MicroArray Profile plugin (OptiNav Inc., Bellevue, WA, USA) for ImageJ v1.47 (http://rsbweb.nih.gov/ii/). Two to four replicates per conditions were obtained per experiment, and all data presented represents means ± s.e.m. from at least three independent experiments.

#### Immunofluorescence

Permeabilized cells were incubated with 200μl of Odyssey Blocking buffer (LI-COR) for 30min. Primary antibodies were applied for 1h and fluorescently tagged (Alexa568) secondary antibodies for 30min, all antibodies were diluted in Odyssey Blocking buffer (LI-COR). Cells were washed three times with PBS between each antibody incubation step. Cells were imaged in PBS at 37°C.

#### mRNA profiling

MCF7 cells were trypsinized and 6×10^5^ cells were suspended in 4ml RNAse free water (Thermo Scientific) with 1ml RNAlater (Thermo Scientific). mRNA extraction and profiling was performed by Comprehensive Biomarker Center GmbH, Heidelberg on an array designed by Agilent 60-mer Sure print technology. The mRNA levels were obtained from three independent runs.

#### Quantifying ectopic EGFR-mTFP expression in MCF7 cells

MCF7 and MCF10A cells were seeded at ~3×10^4^ per well in 8-well Lab-Tek chambers (Nunc). MCF7 cells were transfected with EGFR-mTFP as described previously (see Cell culture and transfection). After serum starvation for 6h, cells were washed once with PBS and treated with EGF-Alexa647 (100ng/ml) for 5min at 37°C. After stimulation, cells were fixed with Roti^®^ Histofix 4% for 10min and their nuclei stained with Hoechst (1μg/ml in TBS) for 5min. Cells were imaged in TBS on a Leica TCS SP8 confocal microscope. The mean EGFR-mTFP and EGF-Alexa647 fluorescence intensity per cell was obtained after cell segmentation in CellProfiler (Kamentsky et al., 2011) using the fluorescence of the nuclear stain (Hoechst) and the EGF-Alexa647. The histograms (Kernel density distribution) obtained from single-cell mean EGF-Alexa647 intensities are shown in Fig.1a.

#### Hydrogen peroxide measurements

Intracellular H_2_O_2_ levels were determined by PF6-AM (Dickinson et al., 2011) (kindly provided by Prof Christopher J. Chang, University of California, Berkeley) fluorescence. MCF7 cells were seeded on 4-well Lab-Tek dishes. The next day, cells were transfected with EGFR-mCherry expression plasmid as described in Transfection above. After starvation in DMEM containing 0.5% FCS for 5-6h, cells were loaded with 5μM PF6-AM in DMEM for 30min at 37°C with or without 320ng/ml EGF-Alexa647. For NOX inhibition, cells were incubated with 10μM DPI 20min before PF6-AM loading. The cells were then washed twice with fresh DMEM and imaged immediately in DMEM (with 25mM HEPES, without Phenol Red) on a Leica TCS SP8 confocal microscope.

#### Temporal H_2_O_2_ profiles upon EGF stimulation

MCF7 cells were transfected with EGFR, cHyPer3 and C1-mCherry (Clontech) as described previously (see Cell culture and transfection). Cells were starved in DMEM containing 0.5% FCS for 5-6h and the medium was exchanged to Hank’s Balanced Salt Solution (HBSS) supplemented with 20mM HEPES. Images were acquired at 1min interval for 20mins on a Leica TCS SP8 confocal microscope. EGF-Alexa647 was added at 5min to a final concentration of 320ng/ml.

#### Detection of PTP_X_ catalytic cysteine oxidation

MCF7 cells were seeded at ~3×10^5^ cells/well in a 6-well culture dish (Nunc) and transfected with 1μg PTP_X_-mCitrine and 1μg EGFR per well. Prior stimulation cells were starved for 6h in supplemented DMEM with 0.5% FCS, followed by treatment with 25mM Dimedone (Sigma-Aldrich) for 10min at 37°C together with EGF-Alexa647 or H_2_O_2_ in serum-free medium. For NOX inhibition, cells were incubated with 10μM DPI for 20min at 37°C prior to Dimedone treatment. After incubation, cells were washed in ice-cold PBS supplemented with 100mM N-Ethylmaleimide (NEM, Sigma-Aldrich) and lysed in 85μL ice-cold lysis buffer (50mM Tris-HCl, pH 7.9, 150mM NaCl, 1% IGEPAL, 0.5% Na deoxycholate, 20mM NEM and protease inhibitors). For immunoprecipitation, equal amounts of protein lysates were incubated with Dynabeads^®^ Protein G magnetic beads (ThermoFisher) and subsequently incubated with anti-GFP antibody overnight at 4°C. Lysates were incubated for 2h with Dynabeads^®^ Protein G for pull down. Total and immunoprecipitated (IP) proteins were resolved by SDS/PAGE using NuPAGE Novex 4-12% Bis-Tris gels (ThermoFisher) in MOPS running buffer, transferred to PVDF membrane and then blocked with LI-COR blocking buffer (LI-COR Biosciences) for 1h. The membrane was then incubated with Anti-Sulfenic acid modified cysteine antibody (Seo and Carroll, 2009) and anti-GFP antibody overnight at 4°C. Next, the membrane was washed with TBS/T and incubated with the respective secondary antibodies for 1h. After washing with TBS/T, the blot was scanned using an Odyssey Infrared Imaging System (LI-COR). Western blot (WB) images were analyzed using FIJI (https://fiii.sc/) and Igor Pro 6.37 (https://www.igorpro.net/). For the temporal cysteine oxidation profiles, MCF-7 cells were stimulated with 5P-EGF in supplemented DMEM. Cells were incubated with 25mM Dimedone 10min before stopping the reaction by ice-cold PBS.

#### Widefield anisotropy

Anisotropy microscopy was performed on an Olympus IX81 inverted microscope (Olympus Life Science) equipped with a MT20 illumination system and a temperature controlled CO2 incubation chamber at 37°C and 5% CO_2_. A linear dichroic polarizer (Meadowlark Optics) was implemented in the illumination path of the microscope and two identical polarizers were placed in an external filter wheel at orientations parallel and perpendicular to the polarization of the excitation light. Fluorescence images were collected via a 20x/0.75 NA air objective using an Orca CCD camera (Hamamatsu Photonics). BFP fluorescence emission was detected between 420-460 nm, mCitrine fluorescence emission between 495-540 nm and Alexa647 fluorescence emission between 705-745 nm.

For each field of view two images were acquired, one with the emission polarizer oriented parallel to the excitation polarizer (*I*_║_) and one with the emission polarizer oriented perpendicular to the excitation polarizer (*I*_⊥_). Fluorescence anisotropy (*r^i^*) was calculated in each pixel *i* by:

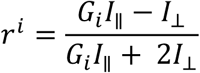

The G-factor (G_i_) was determined by acquiring the ratio of the parallel and perpendicular intensities of Fluorescein in a solution with a steady-state anisotropy close to zero. The CellR software supplied by the microscope manufacturer controlled data acquisition. Live cells were imaged in vitamin-free media in MatTeks and stimulated with EGF-Alexa647. Images were background-substracted and masks of the PM of single cells were generated from the EGFR images using FIJI (https://fiji.sc/).

#### FLIM

Cell arrays were imaged by automated microscopy as described previously (Grecco et al., 2010). An Olympus IX81 microscope (Olympus Life Science) was adapted for frequency domain FLIM. Samples were excited by an Argon laser (Coherent Innova 300C), externally modulated at 79.2MHz through an acousto-optic modulator (AOM, Intra Action SWM-804AE1-1) and fluorescence emission was recorded by a modulated intensified CCD camera (LaVision PicoStar HR / LaVision Imager QE). Both, AOM and image intensifier were modulated with coupled frequency generators (National Instruments PXI-5404). Image stacks were recorded in permuted phase order to reduce bleaching artefacts in the calculation of phase and modulation (van Munster and Gadella Jr., 2004). The setup was controlled by a program developed in-house using LabVIEW 2010 (National Instruments). Phase and modulation were calibrated with a reflective aluminum foil located at the sample plane and drift-corrected with a mirror mounted in a filter cube.

Each cell array microscopy experiment comprised four arrays (for the four different time points: 0, 5, 30 and 120min) glued to a sample holder. The coordinates of the transfection spots were calibrated by automatic localization of six inked reference spots in transmission microscopy with a low magnification objective (UPlanSApo 4x/0.16 NA). To optimize the recording of the number of cells per spot, the array was prescanned in the mCitrine channel with a UPlanApo 10x/0.4 NA objective. The screening then proceeded in two runs with a UPlanSApo 40x/0.9 NA objective, first to obtain the donor-only fluorescence lifetime, followed by a second run after a 4h incubation period with PY72-Cy3.5 to obtain the FRET-FLIM dataset.

The high-content FLIM screening experiments were performed similarly, but the positions were not selected automatically. Here, 2-4 positions in each well were defined and 16-25 fields of view around the selected coordinates were scanned to obtain data from a large number of cells. The complex Fourier components were computed from the phase stack using singular value decomposition. All the data acquired for the same donor molecule (EGFR-EGFP/EGFR-mTFP) and the same batch of labelled antibody (PY72-Cy3.5) was pooled together and jointly analyzed by global analysis (Grecco et al., 2010).

Confocal FLIM experiments to measure EGFR-PTP interactions were performed using a time-correlated single-photon counting module (LSM Upgrade Kit, PicoQuant) on an Olympus FV1000 confocal microscope (see: Confocal microscopy). Pulsed lasers were controlled with the Sepia II software (PicoQuant) at a pulse repetition frequency of 40MHz. The sample was excited using a 440nm diode laser (LDH 440, PicoQuant). Fluorescence emission was spectrally filtered using a narrow-band emission filter (HQ 480/20, Chroma). Photons were detected using a single-photon counting avalanche photodiode (PDM Series, MPD, PicoQuant) and timed using a single-photon counting module (PicoHarp 300, PicoQuant).

Confocal FLIM experiments to measure EGFR phosphorylation were performed on a Leica SP8 confocal microscope equipped with a pulsed 470-670 nm white light laser (white light laser Kit WLL2, NKT Photonics) (see: Confocal microscopy) at 514 nm with a pulse frequency of 20 MHz and emission was restricted with an Acousto-Optical Beam Splitter (AOBS) to 525-550nm. MCF7 cells transfected with EGFR-mCitrine, PTB-mCherry and cCbl-BFP were pulsed for 5min with EGF-Alexa647 (200ng/ml) using the CellASIC ONIX Microfluidic Platform (Millipore) followed by a washout. FLIM measurements were performed prior to and after 5min of EGF stimulation, as well as every 5min after EGF washout for a total of 30min.

For all the confocal FLIM experiments, SymPhoTime software V5.13 (PicoQuant) was used to obtain images after an integration time of 2-4min, collecting app. ~ 3.0–5.0×10^6^ photons per image. For each pixel, the single photon arrival times of the TCSPC measurement were used to calculate the complex Fourier coefficients of the first harmonic and were corrected by the Fourier coefficient of a calculated reference (Grecco et al., 2010).

#### Global analysis of FLIM data

The Fourier coefficients obtained from the FLIM datasets were analyzed by global analysis as previously described (Grecco et al., 2010). Briefly, the global fluorescence lifetimes of the donor alone (*τ*_D_) and donor paired with acceptor (*τ*_DA_) were calculated from the intersection of a linear fit through the Fourier coefficients determined at each pixel with the semicircle corresponding to monoexponential decays. For each pixel, the local fraction of donor molecules that exhibit FRET (α) was calculated from the projection onto the fitted line (Fig. S2a).

#### Confocal microscopy

Confocal images were recorded using an Olympus FluoView FV1000 confocal microscope or a Leica SP8 confocal microscope (Leica Microsystems). The Olympus FluoView FV1000 confocal microscope was equipped with a temperature controlled CO2 incubation chamber at 37°C and a 60x/1.35 NA Oil UPLSApo objective (Olympus Life Science). Fluorescent fusion proteins with BFP, mTFP and mCitrine were excited using the 405nm Diode-UV laser (FV5-LD05, Hatagaya) and the 458/488nm lines of an Argon-laser (GLG 3135, Showa Optronics). Cy3.5/Alexa568 were excited with a 561nm DPSS laser (85-YCA-020-230, Melles Griot) and Alexa647 was excited with a 633nm He-Ne laser (05LHP-991, Melles Griot). Detection of fluorescence emission was restricted as following: BFP (425-450nm), mTFP (472-502nm), mCitrine (525-555nm), Cy3.5/Alexa568 (572-600nm), Alexa647 (655-755nm). Scanning was performed in frame-by-frame sequential mode with 3x frame averaging and a pinhole of 2.5 airy units.

The Leica TCS SP8 confocal microscope (Leica Microsystems) was equipped with an environment-controlled chamber (Life Imaging Services) maintained at 37°C, an HC PL APO 63x/1.4NA CS2 oil objective and an HC PL APO 63x/1.2NA motCORR CS2 water objective (Leica Microsystems). mCitrine, mCherry and Alexa647 were excited with a 470–670nm white light laser (white light laser Kit WLL2, NKT Photonics) at 514nm, 561nm and 633nm, respectively. mTFP was excited by the 458nm Argon laser line, cHyPer3 and PF6-AM by the 488nm line, while BFP was excited with a 405nm diode laser. Detection of fluorescence emission was restricted with an Acousto-Optical Beam Splitter (AOBS): BFP (425-448nm), mTFP (470-500nm), mCitrine (525-551nm), mCherry (580-620nm), Alexa647 (655-720nm) and cHyPer3 (495-530nm). When the oil objective was used, the pinhole was set to 3.14 airy units and 12-bit images of 512×512 pixels were acquired in frame sequential mode with 2x frame averaging. The water objective was used for live cell EGFR trafficking experiments and the pinhole was adjusted (ranging from 3.44 to 2.27 airy units) for each separate channel to maintain optical sectioning fixed to 2.5um.

#### Imaging EGFR vesicular dynamics

Confocal laser scanning microscopy of live MCF7 cells was done on a Leica SP8 confocal microscope (Leica Microsystems) at 37°C using a 63x/1.2NA water objective in DMEM (with 25mM HEPES, without Phenol Red). A temperature-controlled in-house-developed (Masip et al., 2016) flow-through chamber was used to administer a 5min pulsed 200ng/ml EGF-Alexa647 stimulus with the aid of a neMESYS low-pressure syringe pump (Cetoni GmbH). Media were exchanged with a constant flow rate of 3μl/sec to avoid cell detachment. Sustained 200ng/ml EGF-Alexa647 stimulus was administered in 8-well Lab-Tek dishes by pipetting. Images were acquired for ~120min at 1min time intervals. STMs were calculated as described above. The fraction of liganded EGFR-mCitrine was estimated by the EGF-Alexa647/EGFR-mCitrine ratio normalized to the maximal value, whereas the ligandless EGFR-mCitrine fraction by 1-[EGF-Alexa647/EGFR-mCitrine]. The fraction of phosphorylated EGFR at the PM was estimated using the translocation of PTB-mCherry to the PM-localized EGFR-mCitrine. The following quantity was normalized: (*PTB*_*PM*_/(*PTB*_*T*_–*PTB*_*endo*_)/(*EGFR*_*PM*_/*EGFR*_*T*_), where *PTB*_*endo*_ was estimated from the cytosol by intensity thresholding (1.5*std percentile) and removed from the total PTB pool as it is already bound to the phosphorylated EGFR-mCitine in the endosomes. Similarly, the STMs of phosphorylation were estimated by (*PTB*_*PM*_/*PTB*_*T*_)/(*EGFR*_*PM*_/*EGFR*_*T*_) normalized to the previously estimated phosphorylated PM fraction of EGFR.

#### Multiple EGF pulse experiment

MCF7 cells were transfected with PTPRG, PTPN2 or Control siRNA and subsequently with EGFR-mCitrine, PTB-mCherry and cCbl-BFP. For the Rabb11^aS25N^ experiment, Rabb11^aS25N^–mTFP was transfected additionally, without siRNA transfection, and the flow-through chamber protocol was used as in the single-pulse vesicular dynamics experiment. For the siRNA experiments, the cells were transferred to CellASIC ONIX microfluidic switching plate (M04S-03, Millipore) in complete growth media for 3h followed by serum starvation for at least 6h. An EGF pulse-washout program consisting of a 5min pulse of EGF-Alexa647 (20ng/ml) followed by continual perfusion with serum-free media for 25min was delivered using the CellASIC ONIX microfluidic device. Confocal imaging was performed concurrently during 4 successive EGF pulse-washout programs using the Leica TCS SP8. PM phosphorylated fraction of EGFR-mCitrine was estimated in the same manner as for the single-pulse vesicular dynamics experiment.

### Quantification and Statistical Analysis

#### Single cell segmentation and quantification

Cells were segmented in CellProfiler (Kamentsky et al., 2011) using the image of the nuclear stain (Hoechst) and EGFR-mTFP. All images were background-substracted and corrected for bleed through, and mean values per cell (excluding the nuclear region) from all channels were obtained. To match the images of the FLIM MCP and the highresolution CCD camera, the masks were affine transformed (OpenCV).

#### CA-FLIM identification of PTPs that dephosphorylate EGFR

EGFR phosphorylation and the respective phosphorylation fold change (PFC) upon ectopic expression and knockdown of individual PTP_x_s were calculated as *α*_*PTPx*_/*α*^*ctr*^. The corresponding distributions (α_median, PTPx_, α_median, ctr_) obtained from single cells were subjected to a two-sided Kolmogorov-Smirnov (KS) test (SciPy). In case of ectopic PTP_x_-mCitrine expression, if p < 0.05 in > 50% of the experiments (N=4–8), the mean aR was calculated from all significant experiments, otherwise aR was set to 1.

#### Relative specific PTP_X_-mCitrine activity

The α_median_ of each cell was plotted against the respective PTP_X_–mCitrine mean intensity per cell for each time point (Figure S2D). If the distributions of α_median, PTPx_ and α_median, ctr_ were significantly different (Mann–Whitney U, p<0.05), the data was fitted with an exponential function (*α* = *c* + *A* · *e*^‒*k*·*PTPx*^). For each time point, the cells of the respective control measurement were included in the fit after removing outliers (± 3x median absolute deviation around the median). The control coefficient that reflects the dephosphorylating efficiency of each PTP_X_ was determined from the slope of the exponential function at 0 calculated from –k^*^A, where k is the rate and A is the amplitude. For weak α - PTP_X_-mCitrine intensity dependencies, the control coefficients were determined from the slope of a linear fit.

#### Spatial-temporal maps (STMs)

Cells were masked from the EGFR images using FIJI (https://fiji.sc/), the nuclei were segmented using CellProfiler from the nuclear stain (Hoechst) or cCBL-BFP images. For each pixel within the cell, the distance to the closest PM and nuclear membrane (NM) were calculated to derive a normalized distance r = r_PM_ / (r_PM_ + r_NM_). All pixels were split in 10 intervals according to their normalized distances. For each of the observables (EGFR-mTFP, PTP_X_-mCitrine, pY_1068_-Alexa568, and EGF-Alexa647 fluorescence intensities) or derived quantities (α, pY_1068_-Alexa568/EGFR-mTFP, EGF-Alexa647/ EGFR-mTFP, PFC), the mean value was calculated for each segment, yielding a radial profile for the individual cells. To calculate the radial distribution of EGFR-mTFP phosphorylation at distinct pY_i_ sites, the mean fluorescence per segment of the pY_i_ channel was divided by the corresponding mean EGFR-mTFP fluorescence. With the exception of α and pY_1068_/EGFR-mTFP images, all profiles were divided by the total cell mean and an average radial profile was calculated. The radial profiles from the distinct time points were then combined to yield the corresponding spatial-temporal maps. Cells in which PTP_X_-mCitrine expression levels saturated EGFR dephosphorylation were excluded from the analysis (Figure S3D, explanation below).

The STM of the phosphorylation fold-change (PFC) was calculated by dividing the STM pY_1068_/EGFR-mTFP of the control by the STM pY_i_/EGFR-mTFP for each PTP_X_-mCitrine. The profiles from multiple experiments were averaged and significance was determined using 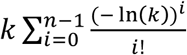, where *k* = Π_*n*_*p_i_* and p_i_ denotes the individual p-values from a Student’s t-test comparing the pY_1068_/EGFR-mTFP distributions of the control to that upon the respective PTP_X_-mCitrine expression at each point in space and time. To obtain the PFC significance, the mean+STD for each STM of pY1068-Alexa568/EGFR-mTFP was calculated per batch, for both the control case and upon PTP_X_ perturbation (cDNA or siRNA). Statistical significance analysis between the two cases was carried out for every spatio-temporal point independently, assuming the data sets are log-normally distributed. Logarithms of the two variables will then give normally distributed variables, which were subtracted using Gaussian addition, effectively calculating the PFC of the batch induced by the PTP_X_ perturbation. For the cDNA case, the control pY1068/EGFR was divided over the respective STM upon PTP_X_-mCitrine expression, whereas for the siRNA case, the ratio was inverted. Using Gaussian product, we then combined the normally distributed variables of the different batches to produce the combined log-PFC. We perform a t-test on this distribution, propagating the degrees of freedom (number of data points), using one-sided Welch's t-test where we checked how statistically significant is the log-PFC distribution relative to zero. To obtain the plots shown in Figure 3B-D, we convert back to a combined log-normal PFC, that can be described though its mean+STD. The spatio-temporal points that are not significantly larger than one (using out previous t-test results of the log-PFC) are shown as transparent in Figure 3B-D.

#### Determining PTP_X_ reactivity towards phosphorylated EGFR

From the EGFR/PTP reaction cycle, the temporal evolution of phosphorylated ligand-bound EGFR (*EGF-EGFR*_*p*_) can be described by:

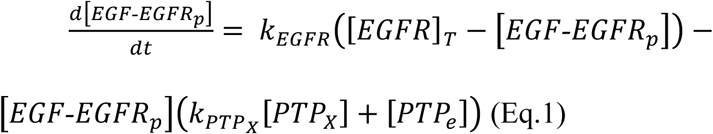

where the forward kinase reaction is assumed to be first order and the backward reaction second order. *k*_*EGFR*_ is the rate constant of kinase activity, *k_PTP_X__* - the PTP_X_-mCitrine dephosphorylation rate constant, [PTP_X_] the concentration of PTP_X_-mCitrine, [EGFR]_T_ the total EGFR concentration and [*PTP*_*e*_] the overall endogenous PTP_e_ activity. Assuming that the local steady-state is reached on a shorter time scale than exchange of EGFR between denoted spatial segments via trafficking (Hendriks et al., 2003), gives:

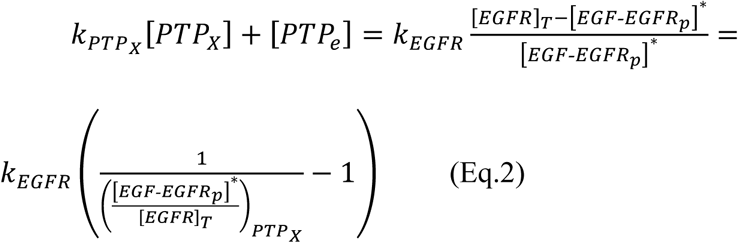

with 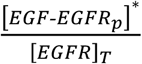 being the fraction of phosphorylated liganded EGFR. In the absence of ectopic PTP_X_-mCitrine expression ([*PTP*_*X*_] = 0), the overall endogenous PTP activity is 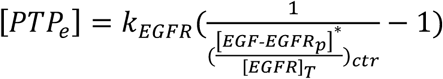, rendering the reactivity of PTP_X_–mCitrine towards specific EGFR-pY_1068_ to be:

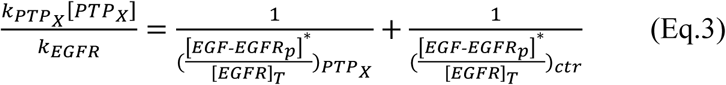

for each spatial-temporal bin. Additionally,

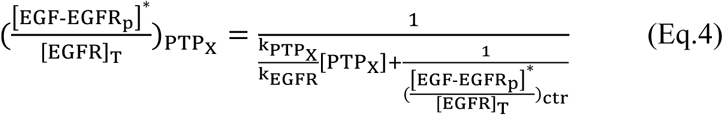

depicts the dependency of the fraction of phosphorylated EGFR on the PTP_X_-mCitrine expression level in steady state and was used to determine PTP_X_-mCitrine expression levels where EGFR phosphorylation was sensitive to the perturbation (Figure S3D).

#### Ligandless EGFR recycling rates

To determine the recycling dynamics of ligandless EGFR upon 5min pulsed EGF stimulus, we developed a dual-compartment model where EGFR internalization from the PM to RE occurs with rate constant *k_in_* and EGFR recycling back to the PM with rate constant *k_rec_*. During the initial 5min stimulus, the PM fraction of ligandless EGFR (*EGFR_PM_*) relative to the total ligandless concentration (*EGFR_T_*) is reduced due to ligand binding. Assuming no further conversion between ligandless and liganded EGFR occurs after removal of EGF, replenishing ligandless EGFR at the PM takes place in the time span of ~5-30min according to the following dynamics:

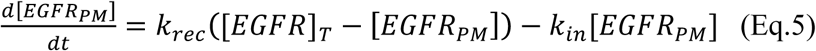

yielding a closed-form solution

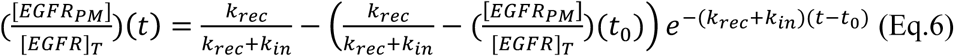

Here, 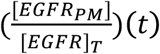 represents the fraction of EGFR at the PM for a particular time *t*, and 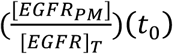 - the PM fraction at t_0_ ≈ 5min. This model was used to infer the trafficking rates from the live cell data, where the first three (out of ten) spatial bins defined the PM (Figure 1G, bottom). Given that in steady state 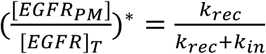 renders

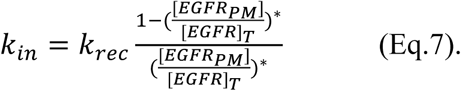

Thus, the steady state PM fraction of ligandless EGFR estimated from the kin vs krec correlation scatter plot (Figure S1K) was ~0.3 with 95% confidence bounds (0.285, 0.311). The estimated average quantities (with 95% confidence bounds) were: k_in_ = 0.325min^-1^ (0.166, 0.483), k_rec_=0.15min^-1^ (0.05, 0.25), and the recycling half-life 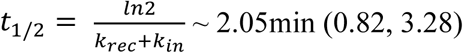.

#### Live cell dose response imaging and quantification

Confocal laser scanning microscopy on live MCF7 cells was performed on a Leica SP8 confocal microscope (Leica Microsystems) using a 63x/1.4NA oil objective. The samples were maintained at 37°C in DMEM (with 25mM HEPES, without Phenol Red). Cells were stimulated every ~1.5min with increasing dose of EGF-Alexa647, ranging from 2.5ng/ml to 600ng/ml (0.34nM to 81.29nM; doses were roughly doubled: D={2.5, 7.2, 16.4, 34.75, 71.6, 145.6, 294.4, 593.4ng/ml}. For NOX inhibition, cells were incubated with 10μM DPI for 30min prior to stimulation. The fluorescence of expressed TagBFP was used to identify the cytoplasmic region of the cell using Otsu’s thresholding method (Otsu, 1979) (scikit-image, scikit-image.org). The PM region of a cell in each time point was calculated by subtracting the cytoplasmic region from the cellular image mask.

PTB–mCherry translocation to (pY_1086_, pY_1148_) PM-bound EGFR-mTFP(mCitrine) for a given EGF-Alexa647 dose *d* ∈ *D* was quantified as:

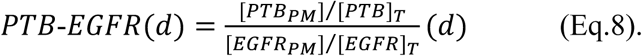

where [PTB_PM_] is the PTB-mCherry translocated to the PM, whereas [PTB]_T_ is the total PTB-mCherry in the cell. The fraction of phosphorylated receptor was then calculated by normalizing this value between the initial (unstimulated) and maximal value of the series

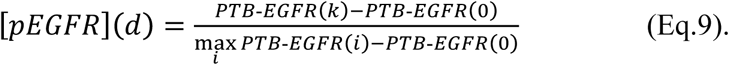

where pEGFR refers to the fraction of phosphorylated EGFR. Similarly, the amount of liganded receptor for dose *d* was calculated from the ratio of integrated EGF-Alexa647 and EGFR-mTFP(mCitrine) fluorescence at the PM:

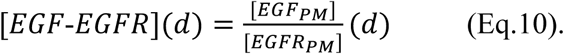

The fraction of liganded receptor (lEGFR) was calculated as:

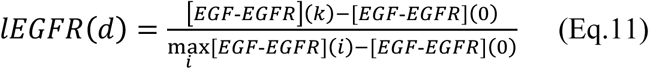

To estimate the relation between the fraction of ligand-bound receptor and the actual administered EGF dose (Figure 1D), the following ligand-binding kinetics model was used:

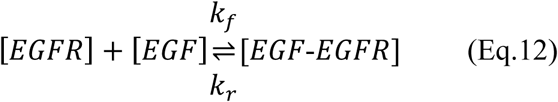

with 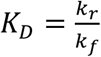 being the dissociation constant. Assuming that at low EGF doses, the ligand will be depleted from the solution due to binding to EGFR (Lauffenburger and Linderman, 1996), the fraction of ligand bound receptor in steady state gives the following closed-form solution:

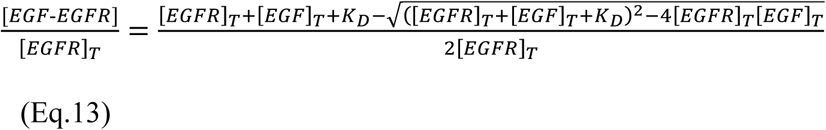

where [*EGFR*]_*T*_ = [*EGFR*] + [*EGF-EGFR*] - the total EGFR concentration on the plasma membrane and [*EGF*]_*T*_ = [*EGF*] + [*EGF-EGFR*] - the total input EGF dose. This function was used to fit the experimental data (Figure S1B,C) thereby mapping the input dose to a fraction of ligand-bound receptor. *K*_*D*_ was obtained to be 760pM.

Area under the curve (AUC) of the dose-response profile of each cell was used as an integrated measure of the response function. The distributions of AUC values between two datasets were compared using two-sample Student’s t-test.

#### Modeling EGFR phosphorylation dynamics

Using the conservation of mass balances:

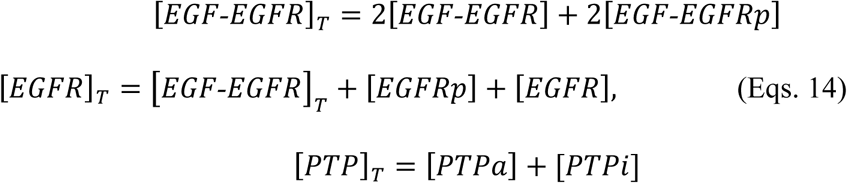

where [*EGF-EGFR*]_*T*_ is the total amount of ligand-bound receptor, [*EGF-EGFR*] - non-phosphorylated ligand-bound EGFR, [*EGF-EGFRp*]–phosphorylated ligand-bound EGFR, [*EGFR*] - ligandless non-phosphorylated EGFR, [*EGFRp*] - ligandless phosphorylated EGFR and [*PTP*]_*T*_ the total amount of ectopically expressed active (*PTP*_*a*_) and inactive (*PTP*_*i*_) PTP. The reaction networks from Figure S4A can be therefore described by the generalized model (Eqs. 15):

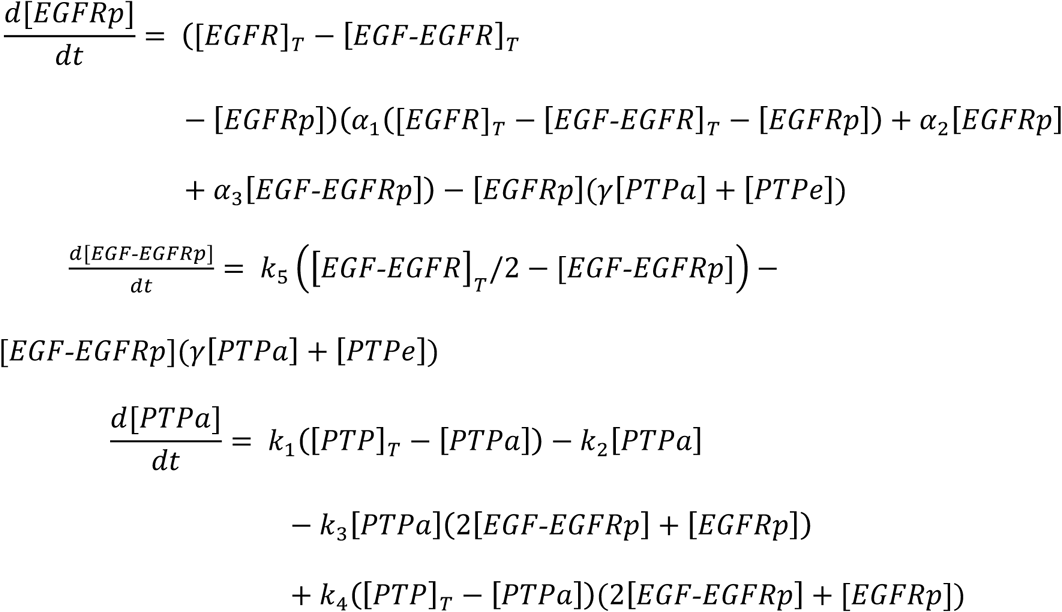

with PTPe contribution given as:

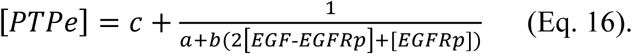

To describe the aggregated effect of endogenous PTP activity on EGFR phosphorylation, the quantities describing ectopic PTP_X_ expression are set to 0, and a, b and c are arbitrary parameters that approximate the aggregated activity of multiple endogenously expressed PTPs. This overall activity was modelled as a combination of double-negative and negative feedback topology as well as negative regulation motifs. In case of ectopic PTP_X_-mCitrine expression on the other hand (*γ* > 0), dephosphorylation of EGFR by PTP_X_-mCitrine will dominate over PTPe, therefore allowing to set [PTP_e_] to zero. Additionally, the following parameter restrictions were imposed: double-negative feedback (k_3_ > 0, k_4_ = 0), negative feedback (k_3_ = 0, k_4_ > 0) or negative regulation (k^3^ = 0, k^4^ = 0).

To determine which of the three EGFR-PTP network topologies (Figure S4A) best represents the experimental EGF dose - EGFR phosphorylation responses upon ectopic PTP_x_-mCitrine expression, the model and data were transformed by expressing the dependency of the fraction of phosphorylated EGFR ([*pEGFR*] = (2[*EGF-EGFRp*] + [*EGFRp*])/[*EGFR*]_*T*_) to the fraction of liganded EGFR-mTFP. The models were then fitted to the data, and the parameters were estimated using an adaptive Metropolis-Hastings algorithm, a variant of the Monte Carlo Markov Chain (MCMC) method for sampling from the posterior joint probability distribution of the parameters (Chib and Greenberg, 1995). Akaike information criterion was used for model discrimination (Hipel, 1981). The parameter values used to fit all EGF-dose EGFR-response curves in Figure 4A-C, and the corresponding Akaike information criterion values are shown in Table S1. The analysis was performed with an in-house developed code in MATLAB (The MathWorks, Inc).

To describe the dynamics of the effective EGFR-PTP network at the PM (Figure 5B,D), the double-negative feedback model (Eqs. 15) was extended with:

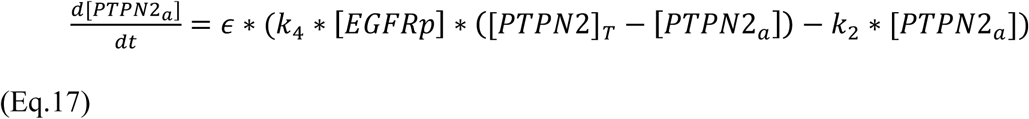

The dephosphorylation of [EGFRp] by PTPN2 was described by an additional term in Eq.15-1: [EGFRp * γ_1_ * [PTPN2_a_]. The EGFR-PTPN2 negative feedback is on a time scale (ϵ) approximately two orders of magnitude slower than the phosphorylation-dephosphorylation reaction, as estimated from the 2-4min recycling time (Figure S1K). The bifurcation analysis of this network was performed using the Bifurcation analysis software XPPAUT (www.math.pitt.edu/~bard/xpp/xpp.html) and interpolated in MATLAB to generate 3D diagrams shown in Figure 5C-D.

## Supplemental item titles

**Table S1 (separate file). Model parameters, Related to Figures 4, 5 and Figure S4.**

**Table S2 (separate file). cDNA-mCitrine expression plasmid constructs, Related to Figures 1-5.** Details of wild type PTP_X_-mCitrine expression plasmids and of mutant PTP_X_-mCitrine and EGFRY845F-mCitrine cDNA expression plasmids.

**Table S3 (separate file). PTP_X_ siRNA SMARTpool, Related to Figure 2.** List of the PTPX with gene name, gene ID, accession no., sequence, pool no. and cat. no. of individual siRNA in a SMARTpool of the human ON-TARGET plus phosphatases siRNA library (Dharmacon).

**Movie S1. EGFR trafficking dynamics upon sustained EGF stimulus, Related to Figure 1H (top).** Composite video comprising of: EGFR-mCitrine (left), EGF-Alexa647 (middle) and EGFR-mCitrine (magenta) – EGF-Alexa647 (green) overlay. Images were taken at 1min interval for ~120min after 200ng/ml EGF-Alexa647 stimulation. Scale bars: 10μm.

**Movie S2. EGFR trafficking dynamics upon 5min pulsed EGF stimulus, Related to Figure 1H (bottom).** Composite video comprising of: EGFR-mCitrine (left), EGF-Alexa647 (middle) and EGFR-mCitrine (magenta) – EGF-Alexa647 (green) overlay. Images were taken at 1min interval for ~120min after 200ng/ml EGF-Alexa647 stimulation. Scale bars: 10μm.

